# The ciliary targeting of membrane proteins by a ternary complex comprising transportin1, Rab8 and the ciliary targeting signal

**DOI:** 10.1101/059451

**Authors:** Viswanadh Madugula, Lei Lu

## Abstract

The sensory functions of cilia are dependent on the enrichment of ciliary resident proteins. While it is known that ciliary targeting signals (CTSs) specifically target ciliary proteins to cilia, it is still unclear how CTSs facilitate the entry and retention of ciliary residents at the molecular level. We found that non-ciliary membrane reporters can passively diffuse to cilia via the lateral transport pathway and the translocation of membrane reporters through the ciliary diffusion barrier is facilitated by importin binding motifs/domains. Screening known CTSs of ciliary membrane residents uncovered that fibrocystin, photoreceptor retinol dehydrogenase, rhodopsin and retinitis pigmentosa 2 interact with transportin1 (TNPO1) via previously identified CTSs. We further discovered that a novel ternary complex, comprising TNPO1, Rab8 and CTS, can assemble or disassemble under the guanine nucleotide exchange of Rab8. Our study suggests a novel mechanism in which TNPO1/Rab8/CTS complex mediates selective entry and retention of cargos within cilia.

## Introduction

Primary cilia (hereafter cilia) are hair-like organelles on the cell surface that can sense diverse environmental cues and initiate corresponding intracellular signaling. Therefore, cilia play important roles in tissue development and homeostasis and defects of cilia can cause a broad range of human genetic diseases, which are collectively called ciliopathies (Nachury et al., 2010). The sensory functions of cilia rely on the presence of a battery of membrane proteins/receptors on the ciliary membrane. How ciliary resident membrane proteins are specifically targeted to cilia is a fundamental question that remains open. While soluble cargos access the ciliary interior from the cytosol via a soluble diffusion barrier at the opening near the cilium base, membrane cargos can follow the following two pathways(Nachury et al., 2010). In the polarized exocytosis pathway, a membrane protein is first packed into a vesicle derived from either the secretory or endocytic pathway. Then the vesicle specifically fuses to the plasma membrane (PM) near the ciliary base (periciliary membrane) and the membrane cargo subsequently enters cilia (Papermaster et al., 1985). In contrast, in the lateral transport pathway, cargos at the PM can directly slide through the ciliary opening to the ciliary membrane without membrane fission or fusion (Hunnicutt *et al.*, 1990; Milenkovic *et al.*, 2009; Leaf and Von Zastrow, 2015). In both pathways, cargos must cross a membrane diffusion barrier at or near the transition zone before entering ciliary membrane(Nachury et al., 2010). The physical and functional existence of such barrier has been substantiated by morphological data from the electron microscopy (EM) (Gilula and Satir, 1972) and kinetic data from the fluorescence recovery after photobleaching (FRAP) (Hu *et al.*, 2010; Chih *et al.*, 2012; Leaf and Von Zastrow, 2015). At molecular level, nucleoporins (Kee et al., 2012), B9 complex (Williams *et al.*, 2011; Chih *et al.*, 2012), Septin2 (Hu et al., 2010) and densely packed membrane lipids (Vieira et al., 2006) have been proposed to contribute to the membrane diffusion barrier. However, it is still unclear how membrane cargos selectively cross the barrier and are retained within cilia.

The targeting of proteins to distinct sub-cellular compartments is mediated by signals which usually comprise linear and short stretches of amino acids. More than a dozen ciliary targeting signals (CTSs) have been discovered although the molecular mechanism behind their targeting is still unknown(Nachury et al., 2010). Available data demonstrate that CTSs neither converge to sequence consensus nor share trafficking machinery (Nachury et al., 2010). Recent discoveries revealed an unexpected role of nucleocytoplasmic transport machinery, especially importing-β1 and transportin1/importin-β2/TNPO1, in the ciliary targeting of an isoform of Crumbs3, KIF17 and retinitis pigmentosa 2 (RP2) (Fan *et al.*, 2007; Dishinger *et al.*, 2010; Fan *et al.*, 2011; Hurd *et al.*, 2011; Kee *et al.*, 2012). Both importing-β1 and TNPO1 belong to β-karyopherin family and are evolutionarily conserved cargo receptors for nucleocytoplasmic trafficking (Marfori et al., 2011). In nucleocytoplasmic trafficking, the weak and transient interactions between importins and FG-repeats of nucleoporins facilitate the importin/cargo complex to cross the diffusion barrier, which is imposed by FG-repeats within the nuclear pore complex (Stewart, 2007; Marfori *et al.*, 2011). In this study, we attempted to elucidate the molecular and cellular mechanism of CTSs and their cognate transport machinery for ciliary membrane proteins. We discovered that a novel ternary complex, comprising TNPO1, Rab8, and CTS, can assemble and subsequently disassemble to transport ciliary membrane cargos under the regulation of Rab8 guanine nucleotide exchange factors (GEFs).

## Results

### Quantification of ciliary localization by the cilium to the PM intensity ratio

We established an image-based and ensemble-averaged metric, the cilium to the PM intensity ratio (CPIR), to quantify the ciliary localization of a membrane protein in cultured mammalian cells. To that end, a line with a width of ~1 μm is drawn orthogonally across the cilium and the maximum of the line intensity profile (I_max_) is subsequently obtained (Fig. 1A-B). After acquiring the mean intensity of the PM (IPM) and the background (I_background_), the CPIR of the membrane protein is defined as 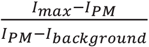.

The CPIR indicates relative enrichment of a membrane protein at the cilium by normalizing its expression level at the PM. Surface labeling can be applied to reduce the interference with the quantification of I_PM_ due to intracellular signal. Importantly, the CPIR was observed to remain relatively constant within > 10-fold expression levels (Fig. 1C and S1A-J).

### PM-localized membrane proteins are able to diffuse passively to the ciliary membrane

CD8a, a PM-localized type I transmembrane protein, is conventionally assumed to be non-ciliary and has been previously used as a reporter to study ciliary targeting of membrane proteins (Follit et al., 2010; Jin et al., 2010). Surprisingly, our imaging data always showed a significant ciliary localization of CD8a and CD8a-GFP in RPE1, BSC-1 and IMCD3 cells (Fig. 1D-E and S2A). We found that the CPIR of CD8a sharply decreased as its cytosolic molecular weight increased by tagging 0-3 copies of GFP (Fig. S2B-D). CD8a-GFP×2 and ×3 were essentially undetectable at cilia (Fig. S2D), implying that the ciliary membrane diffusion barrier has a cytosolic size exclusion limit of 50-100 kDa (considering CD8a forms a homodimer (Rybakin et al., 2011)), similar to the value observed for soluble proteins (Kee et al., 2012).

We initially thought that there could be an uncharacterized CTS in the cytosolic domain of CD8a. However, when both cytosolic and transmembrane domains of CD8a were swapped with corresponding domains of CD4, a type I transmembrane protein natively expressed in only T cells, a significant ciliary localization of CD8a-CD4 chimera was also observed (Fig. S2E). Furthermore, ciliary localization was also observed for diverse PM proteins which are not expected to be ciliary residents, such as interleukin 2 receptor a subunit (IL2Rα, a type I transmembrane protein), GFP-CAAX (lipid-anchored), Vamp5-GFP (tail-anchored), CD59 (glycosylphosphatidylinositol-or GPI-anchored) and endocytosis defective mutants of Vamp2 and 8 (Miller et al., 2011) (Fig. S2E). Supporting our observation, ciliary localization of GFP-GPI and GFP-CEACAM1 were previously reported in IMCD3 cells (Francis et al., 2011). However, we found that not all PM proteins localized to cilia. When the cytosolic tail of CD8a was replaced by that of furin or sortilin, surface labeling revealed that these chimeras localized at clathrin-coated pits instead of cilia (Fig. S2F-H). It has been reported that actin-binding can retain a membrane protein on the PM and prevent it from entering cilia (Francis et al., 2011). Collectively, our data demonstrate that the ciliary membrane diffusion barrier is leaky and PM proteins, if not restrained by clathrin-coated pits or actin cytoskeleton, can enter ciliary membrane non-selectively.

**Figure 1.**
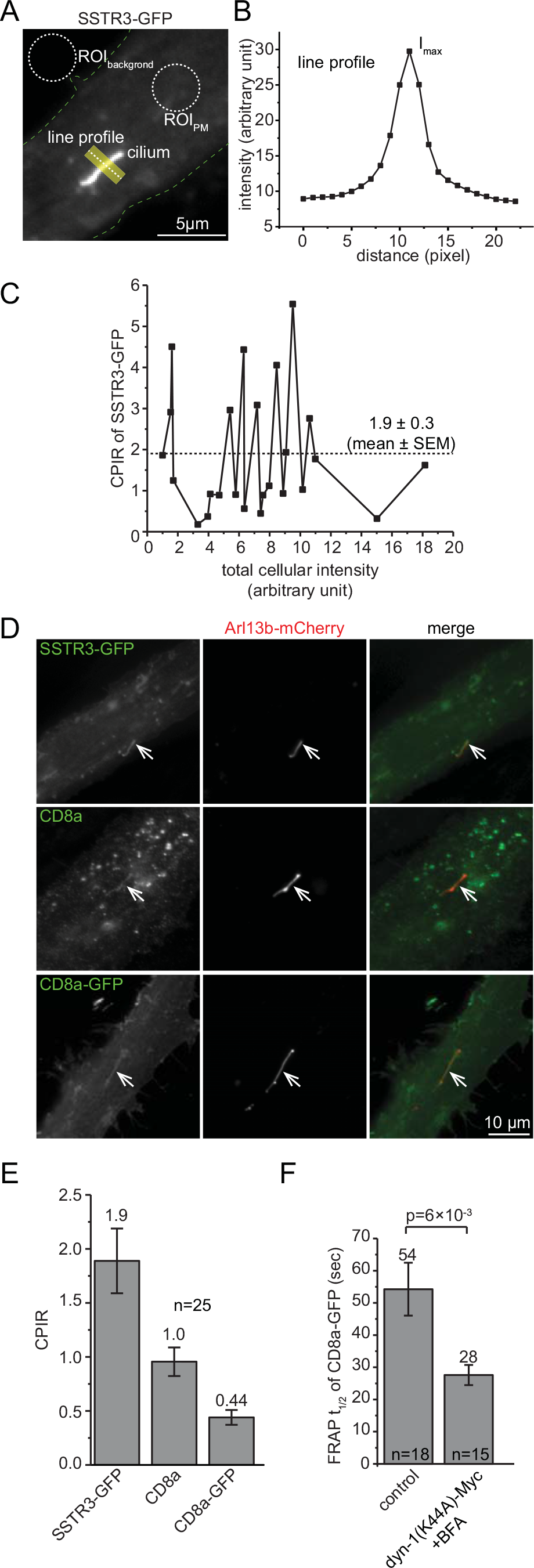
CD8a can access cilia by the lateral transport pathway. (A) A schematic diagram illustrating the acquisition of the CPIR of SSTR3-GFP. A ciliated RPE1 cell expressing SSTR3-GFP was imaged. ROIs of the background (ROL_background_ and PM (ROI_PM_) are shown by dotted circles and used to calculate Ibackground and IPM respectively. The contour of the cell is marked by dotted green lines. A yellow line (with the width of ~1 μm) is drawn across the cilium and the corresponding line intensity profile is shown in (B). Scale bar, 5 μm. (C) Intensity profile of the line. Imax is the peak intensity. (D) The CPIR of SSTR3-GFP is independent on its expression level. For each RPE1 cell expressing SSTR3-GFP, the CPIR is plotted against the total cellular intensity. Data points are connected by lines from low to high total cellular intensity. Dotted horizontal line indicates mean. (D) CD8a and CD8a-GFP are detected in cilia. RPE1 cells transiently co-expressing Arl13b-mCherry and SSTR3-GFP, CD8a or CD8a-GFP were induced for ciliogenesis and imaged. Cilia are indicated by arrows. Scale bar, 10 μm. (E) CPIRs of CD8a and CD8a-GFP. n=25. The mean is indicated at the top of each column. (F) Inhibition of secretory and endocytic pathways did not reduce the recovery kinetics or increase the t_1/2_ during whole cilium FRAP. Ciliated RPE1 cells expressing CD8a-GFP alone (control) or co-expressing CD8a-GFP and dyn-1(K44A)-Myc and treated with BFA were subjected to whole cilium FRAP analysis. The mean and number of cells, n, are indicated at the top and bottom of each column respectively. Error bar, SEM.

We subsequently asked how a PM protein such as CD8a can enter the ciliary membrane. Two pathways are known for the ciliary targeting of membrane cargos: lateral and polarized exocytosis transport pathways. Lacking appropriate sorting signals, CD8a does not undergo polarized secretion or receptor-mediated endocytosis, leaving the lateral transport pathway the most plausible mechanism. Using whole cilium FRAP, we measured the t_1/2_ of CD8a-GFP to be 54 ± 8 sec (mean ± SEM, same for the rest) (n=18) (Fig. 1F and S3A). To rule out the possible contribution of the polarized exocytosis transport pathway, we inhibited the endocytosis by overexpressing dynamin-1(K44A) (van der Bliek et al., 1993) and the secretion by treatment with Brefeldin A (BFA) (Lippincott-Schwartz et al., 2000) (Fig. S3B-C). Under such condition, we found that the t_1/2_ of CD8a-GFP became 28 ± 3 sec (n=15) (Fig. 1F and S3D). Although we do not have a satisfactory explanation for the increased dynamics at the moment, the observation suggests that CD8a and probably other PM proteins could adopt the lateral transport pathway to access the ciliary membrane.

### Importin binding motifs/domains increase ciliary localization of membrane reporters

Since recent studies revealed the role of importins in ciliary targeting (Fan et al., 2007; Hurd et al., 2011), we quantitatively evaluated various importin binding motifs/domains in targeting CD8a to cilia. The cytosolic tail of CD8a was replaced with the following importin binding motifs/domains: 1) the classic nuclear localization signal (cNLS) of SV40 large T antigen, which binds to importin-α and -β1 heterodimer (Marfori et al., 2011), 2) the importin-α binding domain of importin-α (IBB) (Gorlich et al., 1996) and 3) the basic PY-NLS (bPY-NLS) motif of hnRNP-M, which binds to TNPO1 (Lee et al., 2006). The three motifs/domains significantly increased CPIRs of CD8a-GFP derivative reporters (Fig. 2A-D) (p<0.05 by t-test), demonstrating that importin binding motifs/domains can increase the ciliary localization of membrane reporters.

We subsequently employed whole cilium FRAP to study the role of importin binding motifs/domains in ciliary localization of CD8a (Fig. 2E-G). When fused to bPY-NLS or IBB, the CD8a reporter displayed a significantly reduced t_1/2_ — 34 ± 3 sec (n=21) or 29 ± 4 (n=14) respectively, lower than that of CD8a-GFP — 54 ± 8 sec (n=18) (p=0.01 and 0.03 respectively) (Fig. 2F and S3E-F). Their significantly lower t_1/2_s imply the facilitated crossing of the membrane diffusion barrier, consistent with the role of importins in nucleocytoplasmic trafficking (Stewart, 2007; Marfori *et al.*, 2011). Notably, fibrocystin and SSTR3, two ciliary membrane residents, also have lower t_1/2_s than CD8a-GFP (Fig. 2F and S3G-I). However, immobile fractions of both IBB (0.31) and bPY-NLS (0.31) chimeras were similar to that of CD8a-GFP (0.33) (Fig. 2G). In contrast, ciliary membrane residents, such as SSTR3, Arl13b and fibrocystin, displayed much higher immobile fractions (0.63-0.94) (Fig. 2G), as previously reported (Hu et al., 2010; Larkins et al., 2011). Collectively, our data suggest that importins could promote the ciliary localization of membrane proteins by facilitating the ciliary entry instead of retention.

### CTSs of fibrocystin, prRDH and rhodopsin can interact with TNPO1

To test the hypothesis that ciliary membrane residents can utilize importin-β1 or TNPO1 for ciliary targeting, we screened eight CTSs that are known to be sufficient for ciliary targeting. These CTSs, ranging from 7 to 40 residues, were from fibrocystin (Follit et al., 2010), cystin (Tao et al., 2009), polycystin-1/PC-1 (Ward et al., 2011), polycystin-2/PC-2 (Geng et al., 2006), prRDH (Luo et al., 2004), peripherin (Tam et al., 2004), SSTR3 (Jin et al., 2010) and rhodopsin (Tam et al., 2000) (Fig. 3A). GST-fused CTSs of fibrocystin (hereafter f-CTS), prRDH and rhodopsin were observed to specifically pull down endogenous TNPO1 but not importin-β1 (Fig. 3B). Examination of primary sequences of the three CTSs did not reveal PY-NLS consensus motif, which is known to be recognized by TNPO1 (Lee et al., 2006). We first focused on f-CTS for detailed characterization due to its consistently strong interaction with TNPO1.

### f-CTS specifically interacts with TNPO1

Fibrocystin is a type I transmembrane protein of more than 400 kDa, a majority of which is extracellular domain. Lacking the full-length fibrocystin construct, we generated a fibrocystin mimetic fusion protein, named CFF (<u>C</u>D8a luminal domain, <u>f</u>ibrocystin transmembrane domain and <u>f</u>ibrocystin cytosolic domain), by replacing its long luminal domain with corresponding domain of CD8a (Fig. 3C). CFFΔC was generated by deleting the C-terminus of CFF so that it contains only f-CTS in its cytosolic domain, while CD8a-f-CTS was made by replacing the cytosolic domain of CD8a with f-CTS (Fig. 3C). The three chimeras displayed high CPIRs (~8) in ciliated RPE1 cells (Fig. S4A-B). f-CTS-GFP is palmitoylated in cytosol and behaves as a ciliary membrane protein (Follit et al., 2010). When expressed in HEK293T cells, CFF-GFP, CFFAC-GFP, CD8a-f-CTS-GFP and f-CTS-GFP specifically co-immunoprecipitated (co-IPed) endogenous TNPO1 (Fig. 3D-E). The four conserved residues within f-CTS, KTRK, which were reported to be essential for ciliary localization and Rab8 interaction of f-CTS (Follit et al., 2010), were also found to be essential for f-CTS/TNPO1 interaction (Fig. S4B-C). Therefore, we conclude that fibrocystin interacts with TNPO1 via f-CTS. While f-CTS was previously shown to be sufficient for ciliary targeting (Follit et al., 2010), using CFF and its derivatives, we further demonstrated that it is necessary for the targeting. Together with our findings that the CPIR, FRAP t_1/2_ and immobile fraction of CD8a-f-CTS are similar to those of CFF (Fig. 2F-G and S4B), it seems that all ciliary targeting properties of fibrocystin might be attributed to f-CTS.

**Figure 2.**
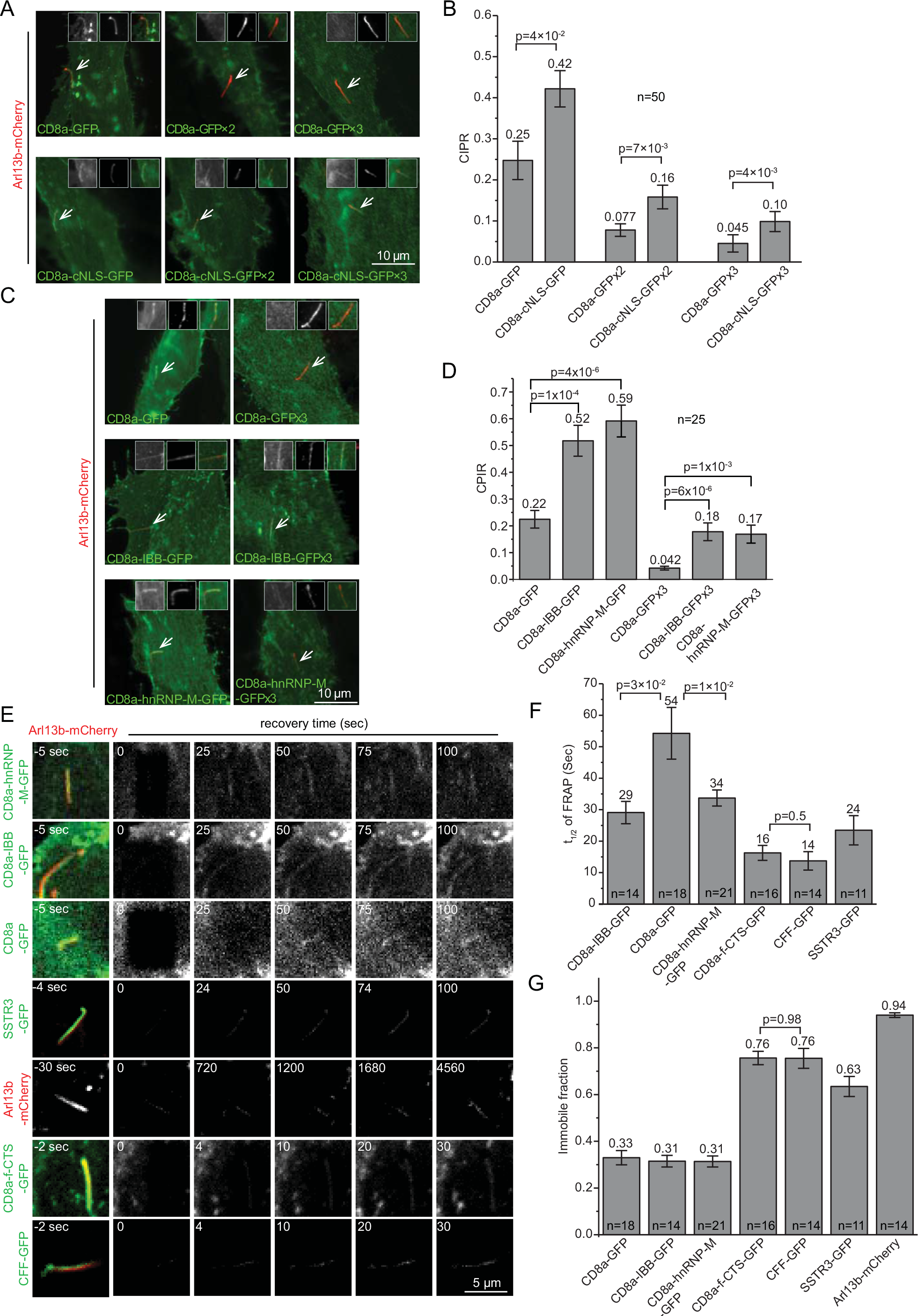
Importin binding motifs/domains increase the ciliary localization of membrane reporters. (A and B) SV40-cNLS increased the ciliary localization of CD8a chimeras tagged with 1-3 GFP molecules. Images showing representative ciliated RPE1 cells co-expressing Arl13b-mCherry and CD8a chimeras. The cilium of interest is indicated by an arrow. The three inserts in each image show the cilium in GFP (monochromatic, left), mCherry (monochromatic, middle) and merge (color, right) channel. Scale bar, 10 μm. The bar graph shows CPIRs of CD8a chimeras. (C and D) IBB and PY-NLS signal of hnRNP-M increased the ciliary localization of CD8a-GFP or-GFP×3. The organization of images and bar graph is similar to (A and B). (E) Representative 2D time-lapse images of cilia expressing various fluorescence chimeras during whole cilium FRAP. Live ciliated RPE1 cells expressing indicated fluorescence chimeras were imaged by a spinning disk confocal microscope. The whole cilium was photobleached at 0 sec. Time is indicated at the upper left of each image. Scale bar, 5 μm. The t_1/2_s and immobile fractions of FRAP are plotted in (F) and (G) respectively. Both CD8a-f-CTS-GFP and CFF-GFP contain the CTS of fibrocystin. Note that the t_1/2_ value of CD8a-GFP in Figure 1E (control) is duplicated here for comparison. The number of cells, n, is labeled in each bar graph. Error bar, SEM. The mean value is indicated at the top of each column. p values (t-test) of selected pairs are denoted.

**Figure 3.**
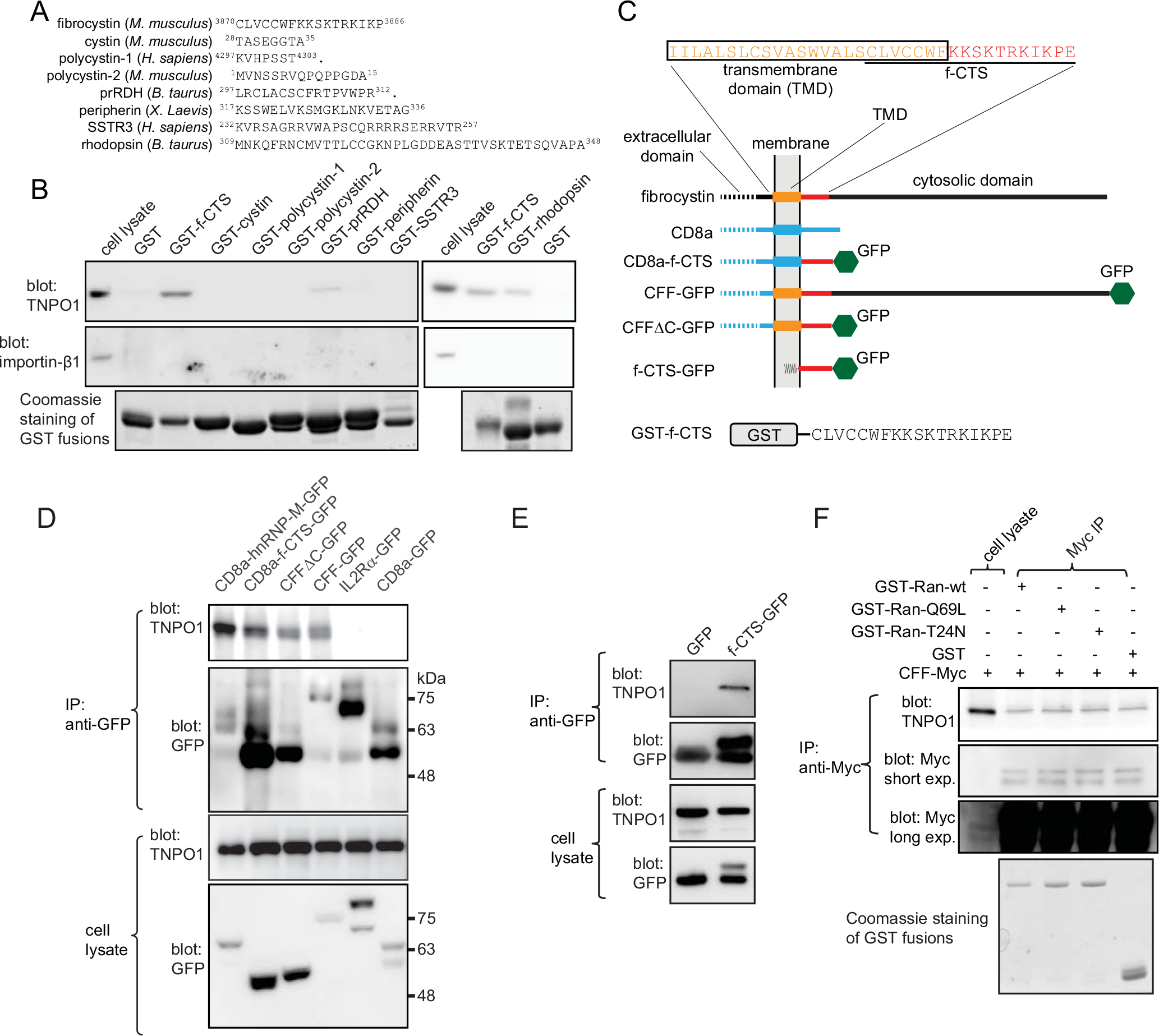
f-CTS interacts with TNPO1. All cell lysates were from HEK293T cells. (A) Sequences of CTSs used in screening. Positions of the first and last amino acids were indicated by numbers. “.” indicates the end of the coding sequence. (B) Screening revealed CTSs of fibrocystin, prRDH and rhodopsin interacted with TNPOI but not importin-β1. Cell lysates were incubated with various bead-immobilized GST-CTSs and the pull-down was blotted forTNPOl and importin-β1. (C) A schematic diagram illustrating various fusion chimeras of fibrocystin used in this study. TMD, transmembrane domain. The amino acid sequences of the TMD and f-CTS are indicated and represented by lines or rectangles of the same color in the diagram. Black and blue lines or rectangles denote corresponding amino acid sequences from fibrocystin and CD8a respectively. Note that f-CTS-GFP attaches the membrane by the palmitoyl group. (D) CD8a chimeras of fibrocystin specifically co-IPed endogenous TNPO1. Cell lysates expressing various GFP-tagged chimeras were subjected to IP using anti-GFP antibody and co-IPed material was blotted by anti-TNPO1 and anti-GFP. CD8a-hnRNP-M-GFP is a positive control. IL2Rα-GFP and CD8a-GFP are negative controls. CD8a chimeras display complex band patterns due to O-glycosylation. In selected gel blots, numbers at the right indicate molecular weight markers in kDa. (E) f-CTS-GFP specifically co-IPed endogenous TNPO1. The co-IP experiment was performed similarly to (D). GFP is a negative control. (F) The interaction between fibrocystin (CFF) and TNPO1 is probably not regulated by Ran GTPase. Cell lysates expressing CFF-Myc were subjected to co-IP using anti-Myc antibody in the presence of the following recombinant proteins: GST (negative control), GST-Ran-wt, -Q69L or -T24N. The co-IPed TNPO1 was subsequently blotted. Two images from short and long exposure (exp.) of the same anti-Myc blot show IPed and cell lysate CFF-Myc respectively.

### The interaction between f-CTS and TNPO1 is probably not regulated by Ran GTPase

Serial truncations further revealed that the region from residue 316 to 539 of TNPO1 is essential for its interaction with f-CTS, while Ran binding region at N-terminus is dispensable (Fig. S4D-E). It is known that the importin/cargo complex can be disassembled by Ran-GTP’s binding to importin (Marfori et al., 2011). Using recombinant GST-Ran wild type (wt), T24N (GDP form) and Q69L (GTP form) mutant, we found that TNPO1 primarily interacted with GST-Ran-Q69L (Fig. S4F), consistent with previous studies (Izaurralde *et al.*, 1997; Siomi *et al.*, 1997; Hurd *et al.*, 2011). However, a saturating amount of GST-Ran-Q69L did not reduce the interaction between CFF and TNPO1 (Fig. 3F). Furthermore, the CPIR of f-CTS-GFP was not affected by overexpressing Ran-Q69L (Fig. S4G-I). Therefore, Ran-GTP probably does not regulate TNPO1 dependent trafficking of fibrocystin, in contrast to its role in the interaction between TNPO1 and RP2 (Hurd et al., 2011). Our data is consistent with previous finding that TNPO1-mediated cargo binding and trafficking can be independent on Ran GTPase (Ribbeck *et al.*, 1999; Lowe *et al.*, 2015).

### TNPO1 is essential for the ciliary targeting of fibrocystin, rhodopsin and prRDH

We quantified the ciliary localization of f-CTS when endogenous importing-β1 or TNPO1 was knocked down. We observed that the depletion of TNPO1 reduced ciliogenesis (Fig. S5A-C). In the remaining ciliated cells, the CPIR of f-CTS-GFP also significantly decreased in comparison to that of the control (Fig. 4A-C and S5D-E). In contrast, the CPIR remained the same as the control upon the depletion of importing (Fig. 4A-C). Similarly, depletion of TNPO1 resulted in significantly reduced CPIRs of full-length rhodopsin and prRDH-CTS (Fig. S5F-H). Therefore, in addition to previously reported RP2 (Hurd et al., 2011), fibrocystin, rhodopsin and prRDH also require TNPO1 for ciliary targeting.

### TNPO1, Rab8 and f-CTS form a ternary complex which is regulated by the guanine nucleotide binding status of Rab8

f-CTS has been reported to preferentially interact with GDP form mutant of Rab8 (Follit et al., 2010). After confirming the interaction (Fig. S6A), we further demonstrated that GST-f-CTS directly interacted with Rab8-GDP using purified His-Rab8-wt, -Q67L (GTP form mutant) and -T22N (GDP form mutant) (Fig. 5A). We subsequently asked how TNPO1, Rab8 and f-CTS interact with each other. To test if Rab8 is necessary for the interaction between f-CTS and TNPO1, we took advantage of our observation that rabbit reticulocyte lysate contains endogenous TNPO1 but not Rab8 (Fig. S6B). The observation that GST-f-CTS pulled down a significant amount of TNPO1 from rabbit reticulocyte lysate (Fig. 5B) demonstrated that the interaction between f-CTS and TNPO1 can be direct and independent of Rab8. To test if Rab8-GDP directly interacts with TNPO1, we used bead-immobilized GST-Rab8 to pull down endogenous TNPO1 in the presence of coexpressed CFF or CD8a and found that Rab8-T22N interacted with TNPO1 only in the presence of CFF-GFP but not CD8a-GFP (Fig. 5C). Similarly, Myc-TNPO1 co-IPed GFP-Rab8-T22N in the presence of CFF-HA but not CD8aAcyto-HA (Fig. S6C). Collectively, our results suggest that f-CTS could simultaneously engage both Rab8-GDP and TNPO1, resulting in the formation of a ternary complex.

To investigate the role of Rab8 in the assembly of this complex, we co-expressed three proteins in cells: HA-TNPO1, CFF-Myc and GFP-Rab8 mutant or GFP (negative control) (Fig. 5D). We found that CFF-Myc co-IPed GFP-Rab8-T22N, but not -Q67L and -wt (Fig. 5D), consistent with our result in Figures 5A and S5A. Although TNPO1 was detected in all co-IPs using CFF-Myc as the bait, ~ 3 folds more TNPO1 was co-IPed in the presence of Rab8-T22N than -Q67L or -wt (Fig. 5D). It seems that endogenous or overexpressed Rab8-wt did not contribute to the binding between f-CTS and TNPO1 under our experimental conditions, therefore retrospectively validated the direct interaction between f-CTS and TNPO1 in our study (Fig. 3 and S4C,E). Although f-CTS can interact with TNPO1 independently of Rab8, our finding that Rab8-GDP instead of -GTP greatly promoted their interaction implies that the ternary complex can be weakened or disassembled by the guanine nucleotide exchange of Rab8 from GDP to GTP.

**Figure 4.**
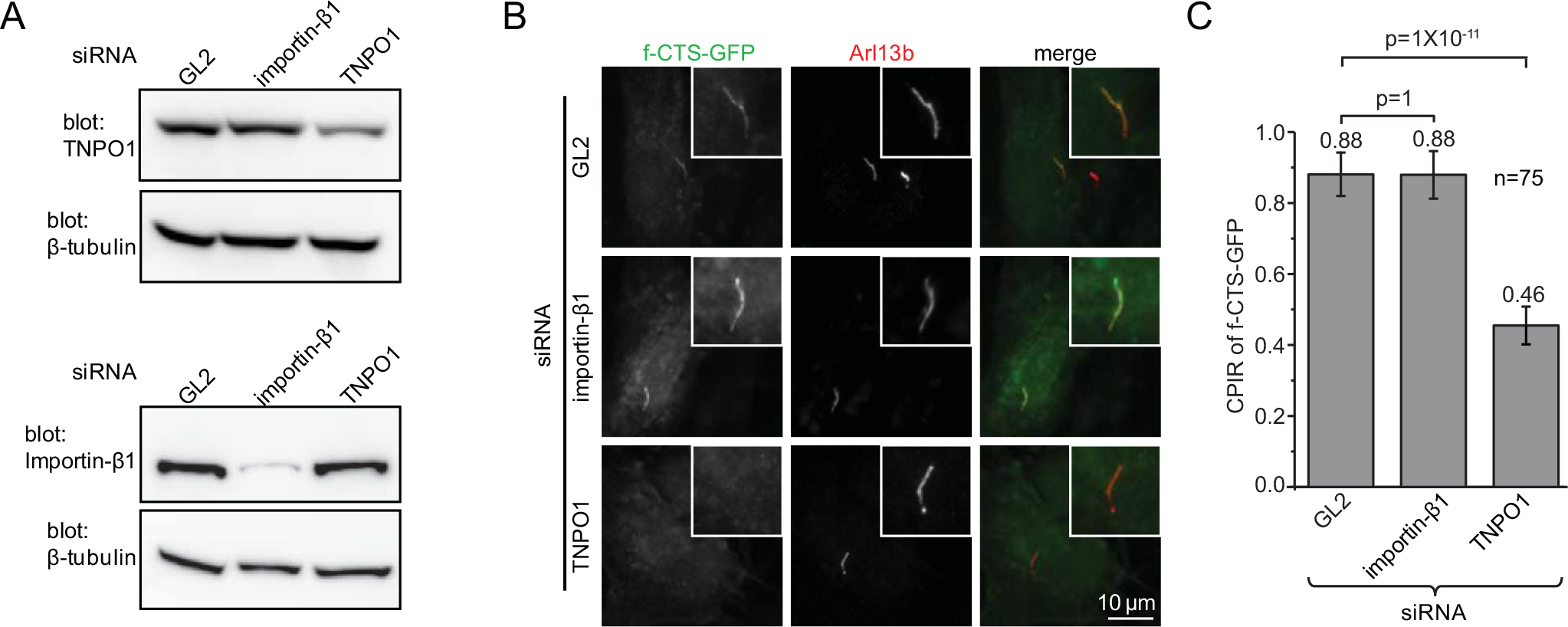
Knockdown ofTNPOl but not importin-βi reduces the ciliary targeting offibrocystin. (A) TNP01 and importin-β1 are specifically knocked down by their siRNAs in RPE1 cells. GL2 siRNA is a negative control. (B and C) RPE1 cells were subjected to knockdown by siRNAs targeting to GL2, importin-β1 and TNP01 respectively. Cells were subsequently transfected to express f-CTS-GFP. Images and CPIRs were acquired after induction of ciliogenesis. In each image, the cilium of interest is enlarged and boxed at the upper right corner. Scale bar, 10 μm. Data in (C) were from n=75 cells. Error bar, SEM. The mean value is indicated at the top of each column. p values (t-test) of selected pairs are denoted.

We next examined the ciliary localization of CFFΔC under the overexpression of Rab8 mutants. As previously reported (Nachury et al., 2007), we observed that Rab8-T22N expression impaired ciliogenesis. In ciliated cells expressing Rab8, the CPIR of CFFΔC decreased significantly in the presence of Rab8-T22N compared to GFP, Rab8-wt and -Q67L (Fig. 5E and S6D), demonstrating the essential role of Rab8 in the ciliary targeting of f-CTS. Since Rab8-T22N is the GDP-locked mutant, GDP to GTP exchange of Rab8 and the ensuing disassembly of the ternary complex is essential for the retention of fibrocystin within cilia. It is possible that, without the guanine nucleotide exchange induced disassembly, the imported TNPO1/Rab8-T22N/f-CTS ternary complex can undergo the reverse pathway, export translocation, to the PM, therefore greatly reducing the CPIR of f-CTS.

### The CTS of prRDH, rhodopsin or RP2 forms similar ternary complex with Rab8 and TNPO1

We wondered if other TNPO1 interacting ciliary membrane residents can assemble similar ternary complex with Rab8 and TNPO1. To that end, we first expanded our study to CTSs of prRDH and rhodopsin which were positive hits in our initial screening (Fig. 3B). The involvement of Rab8 in the ciliary targeting of rhodopsin has been previously reported (Moritz et al., 2001; Wang et al., 2012). Indeed, we found Rab8-T22N but not -Q67L promoted the pull-down of TNPO1 by GST-fused CTS of prRDH or rhodopsin (Fig. 6A-B). We next tested the peripheral membrane protein RP2, which is known to interact with TNPO1 for its ciliary targeting (Hurd et al., 2011). A significantly increased pull-down of TNPO1 by GST-RP2 in the presence of Rab8-T22N was also observed (Fig. 6B). When the major CTS of RP2 was compromised by C86Y and P95L mutations (Hurd et al., 2011), the interaction among RP2, TNPO1 and Rab8-T22N was greatly attenuated (Fig. 6C). Last, we found that RP2-GFP, CD8a-prRDH-CTS-GFP or full-length rhodopsin-GFP was specifically pulled down together with TNPO1 by GST-Rab8-T22N (Fig. S6E), therefore suggesting that RP2, prRDH or rhodopsin could assemble similar ternary complex with Rab8 and TNPO1 via its CTS.

Our finding prompted us to re-examine our initial screen of CTSs in Figure 3B as certain CTS/TNPO1 interactions can take place only in the presence of Rab8-T22N. However, we found that, except CTSs of fibrocystin, prRDH and rhodopsin, the remaining CTSs of our initial screen interacted with neither TNPO1 nor Rab8 in the presence of overexpressed Rab8-T22N (Fig. S6F), suggesting that the utilization of Rab8 and TNPO1 as ciliary transport machinery could be a privilege for certain ciliary membrane proteins.

## Discussion

The PM and ciliary membrane share the same membrane sheet, yet their proteins and lipids do not freely mix due to the membrane diffusion barrier at the ciliary base (Nachury et al., 2010). It is however not understood how ciliary membrane residents cross the membrane diffusion barrier and achieve their retention within cilia. We demonstrated that PM proteins can passively diffuse across the membrane diffusion barrier to cilia, possibly via the lateral transport pathway. Similar to cilia, the inner nuclear membrane (INM) is in direct continuity with the outer nuclear membrane (ONM) and the endoplasmic reticulum (ER) and its distinct composition is maintained by nuclear pore complexes, which function as membrane diffusion barriers between the INM and ONM (Hetzer et al., 2005). Our results on cilia parallel what we know about the INM, as ER membrane proteins can also passively and laterally diffuse to the INM but they have much weaker retention there than *bona fide* INM residents (Zuleger et al., 2012). Two mechanisms have been proposed for the INM targeting: retention and selective entry mechanism (Katta et al., 2014).

**Figure 5.**
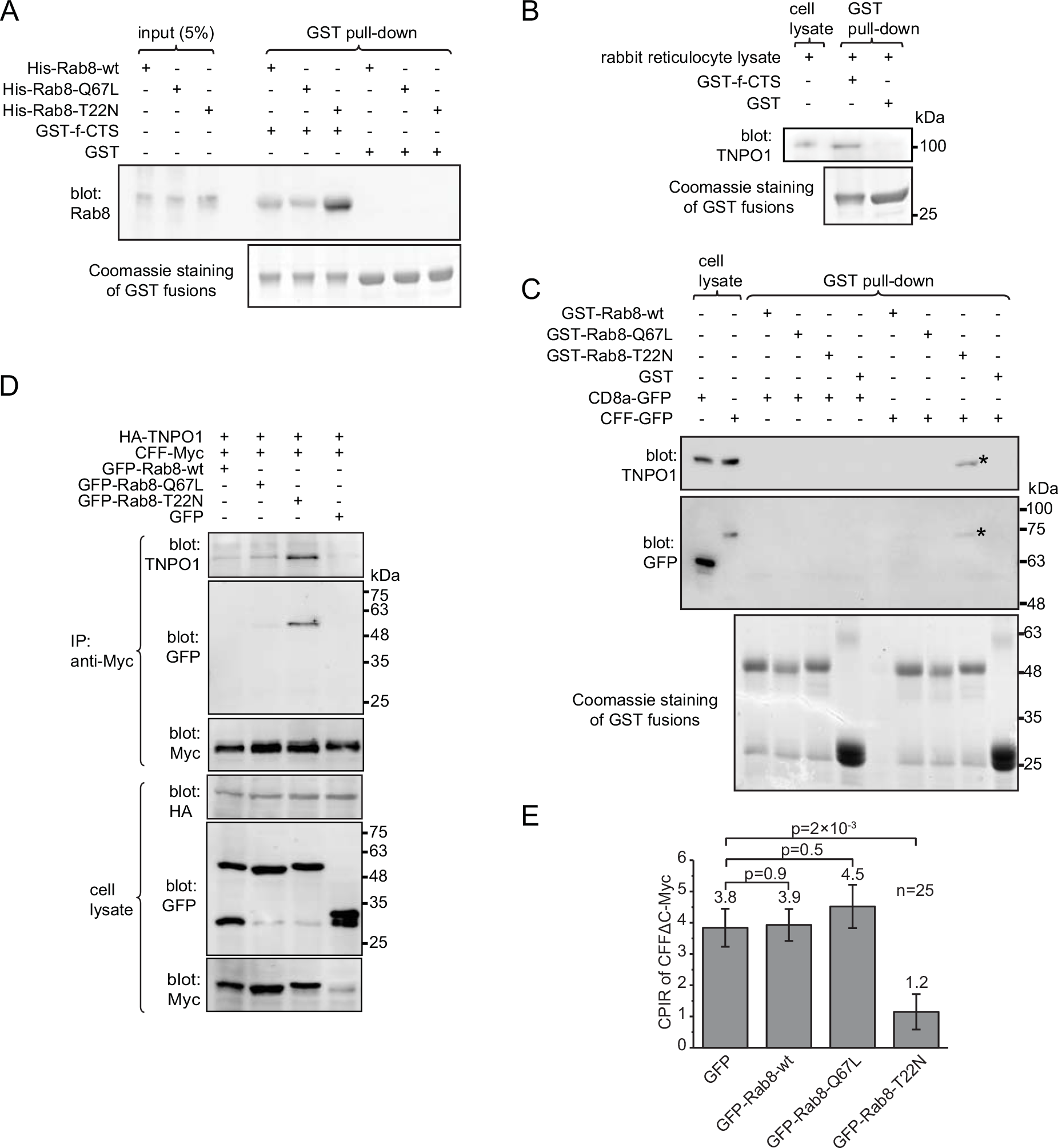
TNPO1, Rab8 and f-CTS form a ternary complex. All cell lysates were from HEK293T cells. (A) f-CTS interacted directly with Rab8-GDP mutant form. Bead-immobilized GST-f-CTS or GST was incubated with purified His-Rab8-wt, -Q69L and -T22N. The protein pulled down was blotted by anti-Rab8 antibody. (B) The interaction between f-CTS and TNPO1 should be direct. Bead-immobilized GST-f-CTS specifically pulled down TNPO1 from rabbit reticulocyte lysate, which contains endogenous TNPO1 but not Rab8 (Fig. S6B). (C) Rab8-T22N interacted with TNPO1 indirectly via fibrocystin. Bead-immobilized GST-Rab8 fusion proteins were incubated with cell lysate expressing CD8a-GFP or CFF-GFP and material pulled down was blotted for TNPO1 and GFP chimeras. “*” indicates the specific protein band. (D) Rab8-T22N promoted the interaction between fibrocystin and TNPO1. The cell lysate triply expressing HA-TNPO1, CFF-Myc and one of the following chimeras, GFP-Rab8-wt, -Q67L, -T22N and GFP, was subjected to IP using anti-Myc antibody and co-IPed TNPO1 and GFP chimeras were blotted. (E) Overexpression of Rab8-GDP mutant reduced the ciliary localization of fibrocystin. CPIRs of CFFΔC-Myc in ciliated RPE1 cells co-expressing CFFΔC-Myc and one of the following chimeras, GFP-Rab8-wt, -Q67L, -T22N and GFP. The mean value is indicated at the top of each column. n=25. Error bar, SEM. p values (t-test) of selected pairs are denoted. In all gel blots, numbers at the right indicate the molecular weight markers in kDa.

**Figure 6.**
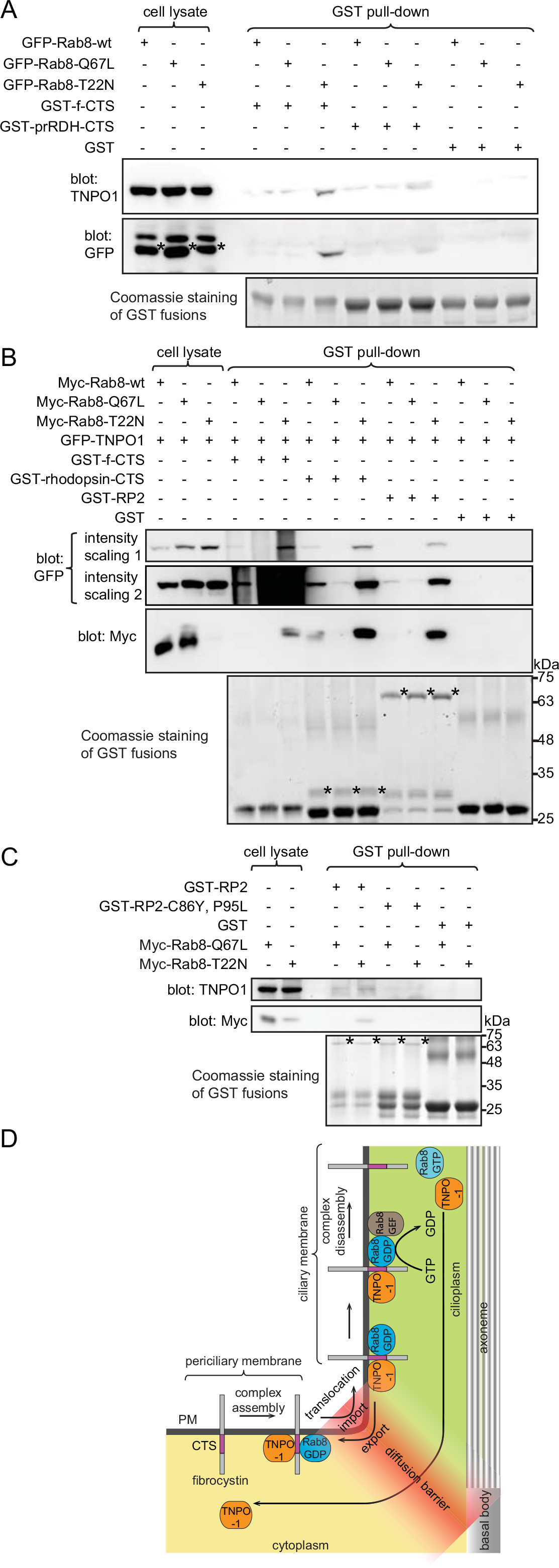
The CTS of RP2, prRDH or rhodopsin can assemble similar ternary complex with Rab8 and TNPO1. (A) Rab8-T22N increased the binding of prRDH-CTS to TNPO1. HEK293T cell lysate expressing GFP-Rab8-wt, -Q67L or -T22N was subjected to pull-down by bead-immobilized GST-prRDH-CTS, GST-f-CTS (positive control) and GST (negative control) and the material pulled down was blotted for GFP-Rab8 and endogenous TNPO1. “*” denotes the band specific to GFP-Rab8. (B) Rab8-T22N increased the binding of rhodopsin-CTS and RP2 to TNPO1. HEK293T cell lysate co-expressing GFP-TNPO1 and Myc-Rab8-wt, -Q67L or -T22N was subjected to pull-down by bead-immobilized GST-f-CTS (positive control), GST-rhodopsin-CTS, GST-RP2 and GST (negative control) and the material pulled down was blotted for Myc-Rab8 and GFP-TNPO1. “*” denotes the corresponding GST-fusion protein used as the bait for the pulldown. (C) The interaction among RP2, Rab8-T22N and TNPO1 is abolished by C86Y and P95L mutation of RP2. HEK293T cell lysate expressing Myc-Rab8-wt, -Q67L or -T22N was subjected to pull-down by bead-immobilized GST-RP2, GST-RP2-C86Y,P95L and GST (negative control) and the material pulled down was blotted for Myc-Rab8 and endogenous TNPO1. “*” denotes the corresponding GST-fusion protein used as the bait for the pull-down. The intensity scaling of the same anti-GFP blot is adjusted to show both bright (intensity scaling 1) and weak bands (intensity scaling 2). In selected gel blots, numbers at the right indicate the molecular weight markers in kDa. (D) Model of ciliary targeting of fibrocystin. See the main text for the description.

CTSs could adopt two similar mechanisms. 1) The retention mechanism prevents ciliary cargos from exiting cilia. The retention can be mediated by the specific binding of CTSs to their cognate ciliary receptors such as BBSome (Jin et al., 2010) and axonemal microtubule (Fan *et al.*, 2004; Kovacs *et al.*, 2008; Francis *et al.*, 2011). Paradoxically, most known ciliary membrane residents are highly mobile within cilia (Hu *et al.*, 2010; Chih *et al.*, 2012; Breslow *et al.*, 2013; Ye *et al.*, 2013). In contrast to ciliary membrane residents, INM proteins are largely immobile (Ellenberg et al., 1997). 2) In the selective entry mechanism, transport receptors selectively facilitate the crossing of ciliary residents through the diffusion barrier by binding to their CTSs. Studies have shown that importins can act as ciliary transport receptors (Fan *et al.*, 2007; Dishinger *et al.*, 2010; Hurd *et al.*, 2011) similar to their roles in the nucleocytoplasmic trafficking (Stewart, 2007; Marfori *et al.*, 2011).

Our FRAP data suggest that importins could promote the selective entry of cilia by facilitating the crossing of the membrane diffusion barrier without increasing the ciliary retention of cargos. We identified four native ciliary membrane residents — fibrocystin, prRDH, rhodopsin and RP2 that specifically form a novel ternary complex with TNPO1 and Rab8-GDP via their CTSs. It has been reported that Rab8-GTP is enriched while-GDP is depleted in cilia (Nachury et al., 2007). The Rab8-GTP gradient which is probably maintained by the polarized ciliary localization of Rab8 GEFs — Rabin8 (Hattula et al., 2002) and RPGR (Murga-Zamalloa et al., 2010). Our findings reveal a novel molecular and cellular role of Rab8 and TNPO1 in non-vesicular ciliary trafficking and the following model is conceivable for the ciliary targeting of membrane cargos such as fibrocystin (Fig. 6D). First, fibrocystin can follow either lateral transport or polarized endocytosis to the periciliary membrane. Next, near the basal body, Rab8-GDP can be released from its GDP dissociation inhibitor (GDI) by Dziβ1, the basal body localized GDI displacement factor for Rab8 (Zhang et al., 2015). The association between f-CTS and Rab8-GDP further recruits TNPO1 to assemble the ternary import complex. Then, facilitated by TNPO1, the complex translocates across the membrane diffusion barrier (selective entry mechanism). After the import translocation, the GDP of Rab8 is exchanged to GTP by cilium-localized GEFs and the ternary complex subsequently disassembles, therefore releasing free fibrocystin to the ciliary membrane. Lastly, with the export of TNPO1, the exit of fibrocystin to the PM is prohibited by the membrane diffusion barrier, hence providing a robust retention mechanism to confine the dynamically diffusive movement of fibrocystin within ciliary membrane.

It is possible that the guanine nucleotide exchange of Rab8 is the rate-limiting step. Consequently, newly imported TNPO1/Rab8/CTS ternary complex can exit to the PM via reversible pathway – export translocation, hence resulting in a rapid but small fraction of recovery during whole cilium FRAP (Fig. 2F-G). Our model bears similarity to the non-vesicular Ran GTPase-dependent nucleocytoplasmic transport pathway, in which Ran-GTP binds to importin to disassemble importin/cargo complex in the nucleus. The Ran-GTP gradient, which is maintained by its nucleus-localized GEF and cytoplasm-localized GTPase-activating protein (GAP), drives nuclear trafficking directionally (Stewart, 2007).

It is tempting to speculate that Rabs and importins could cooperatively target cargos to cilia. Supporting this view, Rab23 and TNPO1 have been reported to assemble a potential complex for the ciliary targeting of KIF17 (Lim and Tang, 2015). Importins possess repeats of domains that can form diverse interfaces to engage a large repertoire of cargos (Marfori et al., 2011). Therefore, more ciliary residents are expected to assemble ternary complexes with Rabs and importins for ciliary targeting. It could be informative to systematically screen ciliary residents for their Rab-dependent interaction with importins.

## Materials and methods

### Knockdown

The following siRNA oligonucleotides were purchased from Dharmacon: GL2 (5’-CGUACGCGGAAUACUUCGA-3’), siRNA smart pool targeting importin-β1 (#L-017523-00-005) (5’-GAACCAAGCUUGAUCUGUU-3’, 5’-GCUCAAACCCCACU AG U U AU A-3’, 5’-G ACG AGAAG UCAAG AACU A-3’, 5’-GGGCGGGAGAUCGAAGACUA-3’) and siRNA smart pool targeting TNPO1 (#L-011308-00-005) (5’-GCAAAGAUGUACUCGUAAG-3’, 5’-G U AU AG AG AU GCAG CC U U A-3’, 5’-GUAAAUACCAGCAUAAGAA-3’ and 5’-GCAAAUGUGUAUCGUGAUG-3’). siRNAs were transfected to RPE1 cells using Lipofectamine 2000 according to manufacturer’s protocol. For the expression of exogenous proteins after knockdown, transfections were conducted after 24 hours of siRNA transfection. 48 hours after the transfection of siRNA, cells were serum-starved for another 48 hours before immunofluorescence labeling.

Endogenous TNPO1 was also depleted by lentivirus-mediated transduction of shRNA. 293FT cells were seeded on 0.01% poly-L-lysine coated 6-well plate. At 60-70% confluency, cells were transfected with packaging plasmids: pLβ1, pLP2, pLP/VSVG (Invitrogen) and lentiviral shRNA construct targeting TNPO1 in the ratio of 2:1:1:4 using Lipofectamine 2000. After 18 hours of transfection, the medium was replaced with fresh one. After 36-48 hours of transfection, the virus-containing medium was collected and filtered to remove cell debris. For shRNA-mediated knockdown of TNPO1, the lentivirus filtrate was immediately used to incubate with RPE1 cells for 12 hours, followed by a second infection with fresh filtrate for another 12 hours. Cells were selected in puromycin to enrich lentivirus infected cells. This stable pool of cells was seeded on coverslips and transfected to express f-CTS-GFP. After the induction of cilia by serum starvation, cells were processed for immunofluorescence labeling.

### Antibodies

The following primary antibodies are commercially available: anti-acetylated α-tubulin (Sigma, #6-11B-1), anti-α-tubulin (Santa Cruz, #sc8035), anti-β-tubulin (Santa Cruz, #sc5274), anti-GAPDH (Santa Cruz, #sc25778), anti-GFP (mouse monoclonal) (Santa Cruz, #sc9996), anti-GFP (rabbit polyclonal) (Santa Cruz, #sc8334), anti-importin-β1 (Abcam, #ab2811), anti-TNPO1 (Abcam, #ab10303), anti-Myc (Santa Cruz, #sc40), anti-CD8a (Developmental Studies Hybridoma Bank, clone OKT8), anti-IL2Ra (ATCC, clone 2A3A1H), anti-Rab8 (BD biosciences, #610844) and HRP-conjugated anti-HA antibody (GeneScript, #A-00169). HRP conjugated goat anti-mouse and -rabbit IgG antibodies were purchased from Bio-Rad. HRP conjugated protein A was purchased from Abcam. Alexa Fluor conjugated goat anti-mouse and -rabbit IgG antibodies were purchased from Invitrogen.

### Cell culture and transfection

hTERT RPE1 and IMCD3 cells were maintained in DMEM and Ham’s F12 mixture medium supplemented with 10% fetal bovine serum at 37°C in 5% CO_2_ incubator. HeLa, BSC-1, HEK293T and 293FT cells were cultured in DMEM High glucose medium supplemented with 10% fetal bovine serum at 37°C in 5% CO_2_ incubator. HeLa and HEK293T cells were transfected using polyethylenimine (Polysciences, Inc.). RPE1, IMCD3, 293FT and BSC-1 cells were transfected using Lipofectamine 2000 (Invitrogen). Transfection was performed when cells reached 70-80% confluency according to standard protocol. To induce ciliogenesis after the overexpression or knockdown of target proteins by transfection, cells were serum starved by incubating in DMEM. Typical starvation time for RPE1, IMCD3 and BSC-1 cells was 2, 2 and 5 days respectively.

### Purification of GST fusion proteins

This was performed as previously described (Mahajan et al., 2013; Zhou et al., 2013). Briefly, DNA plasmids encoding GST fusion proteins were transformed into BL21 *E. coli* cells. Transformed bacteria were induced by Isopropyl P-D-1-thiogalactopyranoside and pelleted. The pellet was resuspended in GST lysis buffer containing 50 mM Tris pH 8, 150 mM NaCl, 5 mM MgCl_2_, 1% Triton X-100, 10 μM DNaseI (Sigma), 0.2 mg/ml lysozyme (Sigma), 1 mM phenylmethylsulfonyl fluoride and 1 mM dithiothreitol and lysed by freeze-thaw method. The lysate was cleared at 13,000×g for 30 minutes and the supernatant was incubated with pre-washed glutathione Sepharose 4B (GE Healthcare Life Sciences) at 4 °C for 2 hours. The beads were washed with the buffer containing 50 mM Tris pH 8, 150 mM NaCl, 1% Triton X-100 and 1mM dithiothreitol. The bead-immobilized GST fusion proteins were subsequently used for pull-down or antibody purification.

### Purification of His-tagged Rab8 fusion proteins

pET30ax DNA plasmids encoding His-tagged Rab8-wt, -T22N and -Q67L were transformed into BL21 *E coli* cells. Transformed bacteria were induced, pelleted and lysed as the purification of GST fusion proteins. The lysate was subjected to centrifugation at 13,000×g for 30 minutes and the supernatant was incubated with pre-washed Ni-NTA agarose beads (QIAGEN) in the presence of 10 mM imidazole at 4°C for 2 hours. After beads were washed with the buffer containing 20 mM HEPES pH7.3, 200 mM KCl, 10% glycerol and 25 mM imidazole, the bound protein was eluted with the elution buffer containing 20 mM HEPES pH7.3, 200 mM KCl, 250 mM imidazole, 10% glycerol and 1 mM DTT. The eluted protein was dialyzed and concentrated.

### Generation of Arl13b polyclonal antibody

His-tagged Arl13b-C-ter was purified under denaturation condition using 8M urea as previously described (Mahajan et al., 2013). The denatured protein was used to immunize rabbits and anti-sera were collected by Genemed Synthesis Inc. To purify the Arl13b antibody from the anti-serum, GST-Arl13b-C-ter on glutathione Sepharose beads were prepared as described above and incubated with dimethyl pimelimidate (Sigma) in 200 mM sodium borate solution pH 9.0 to cross-link the fusion protein to glutathione. After blocking the excess cross-linker by ethanolamine, the cross-linked beads were incubated with anti-serum at room temperature. The beads were subsequently washed with PBS and the bound antibody was eluted by 100 mM glycine pH 2.8. The pH of the eluate was adjusted to neutral immediately and the eluted antibody was dialyzed, concentrated, quantified and stored at-80 °C.

### *in vitro* transcription and translation

The in vitro transcription and translation of Myc-TNPO1 or Myc-Rab8-wt was conducted using TNT^®^ T7 Quick Coupled Transcription and Translation System (Promega) according to manufacturer’s protocol. The reaction mixture was incubated at 30°C for 90 minutes. The protein expression was verified by Western blot analysis.

### Immunoprecipitation and GST pull-down

HEK293T cells were subjected to transfection as described above. After 24-36 hours, cells were scraped in lysis buffer containing 50 mM HEPES pH7.3, 150 mM NaCl and 1% Triton X-100 and the resulting lysate was cleared by centrifugation at 16,000×g at 4°C. The supernatant was incubated with ~1μg of antibody, 15 μl GFP-Trap beads (ChromoTek) or 10-40 μg GST fusion protein on glutathione beads for 4-14 hours. When antibody was used, the antigen-antibody complex was subsequently captured by 15 μl pre-washed Protein A/G beads (Pierce) for 2-4 hours. Beads were washed with lysis buffer and bound proteins were eluted by boiling in SDS-sample buffer and resolved in 8-12% SDS-PAGE. The separated proteins were transferred to polyvinyl difluoride membrane (Bio-Rad). After primary and HRP conjugated secondary antibody incubation, the chemiluminescence signal was detected by a cooled charge-coupled device camera (LAS-4000, GE Healthcare Life Sciences).

### Immunofluorescence microscopy

Cells were seeded on Φ12 mm coverslips (No. 1.5) in a 24-well plate. After 24 hours of transfection, cells were serum starved to induced ciliogenesis and subsequently fixed by 4% paraformaldehyde in PBS at room temperature for 20 minutes. This was followed by neutralizing paraformaldehyde with 100 mM ammonium chloride and washing with PBS. The primary and secondary antibodies were diluted in fluorescence dilution buffer (PBS supplemented with 5% fetal bovine serum and 2% bovine serum albumin) containing 0.1% saponin (Sigma). Cells were incubated with primary antibody, washed and followed by fluorescence-conjugated secondary antibody. After extensive washing, coverslips were mounted in Mowiol 4-88 (EMD Millipore). For surface labeling, cells grown on coverslips were incubated with CD8a monoclonal antibody on ice for 1 hour. After washing with ice-cold PBS, cells were fixed and subjected to immunofluorescence labeling as described above. Cells were imaged under a wide-field microscope system comprising Olympus IX83 equipped with a Plan Apo oil objective lens (63X or 100X, NA 1.40), a motorized stage, motorized filter cubes, an scientific complementary metal oxide semiconductor camera (Neo; Andor,) and a 200 W metal-halide excitation light source (Lumen Pro 200; Prior Scientific,). Dichroic mirrors and filters in filter turrets were optimized for GFP/Alexa Fluor 488, mCherry/Alexa Fluor 594 and Alexa Fluor 647. The microscope system was controlled by MetaMorph software (Molecular Devices) and only center quadrant of the camera sensor was used for imaging.

### FRAP

RPE1 cells were seeded on Χ35 mm glass-bottom Petri-dish (MatTek) and transfected to co-express GFP-fused membrane reporters and Arl13b-mCherry (as a ciliary marker). After 24 hours of transfection, cells were serum-starved to induce cilia. Live cell imaging was conducted in CO_2_-independent medium (Invitrogen). 2D time-lapse images during FRAP were acquired by a motorized Nikon Eclipse Ti microscope equipped with Plan-Apo oil lens (100×, NA 1.4), Perfect Focus System, Piezo z-stage, CSU-22 spinning disk scan head (Yokogawa), 37°C heated chamber (Lab-Tek), 100 mW diode lasers (491 nm and 561nm), 3D FRAP system (iLAS^2^, Roper Scientific) and electron-multiplying charge-coupled device (Evolve 512, Photometrics). The microscope system was controlled by MetaMorph and iLAS^2^ software (Roper Scientific). The 2D time-lapse images of cilia were collected before and after photobleaching. Image analysis was conducted in ImageJ (http://imagej.nih.gov/ij/). The ROI (region of interest) of cilia was generated by either intensity segmentation or manually drawn according to Arl13b-mCherry. The mean fluorescence intensity of ROI at each post-photobleaching time point was fitted to a single exponential decay function y=y_0_+A_1_×*exp(−(x−x_0_)/t_1_) using OriginPro8.5 (OriginLab). The immobile fraction was calculated as (I_pre_−y_0_)/(I_pre_−I_0_), where I_pre_ and I_0_ are the intensity before and immediately after photobleaching.

### Statistical analysis

All samples were randomly chosen and included for analysis. The sample size n is indicated wherever applicable in figures or corresponding figure legends. Data are presented as the mean ± SEM. The two-tailed unpaired t-test was conducted in Excel (Microsoft) and used for the analysis of statistical significance. A p value of less than 0.05 was considered statistically significant.

## Acknowledgements

We would like to thank J. Brenman, P. De Camilli, T Galli, B. Gumbiner, W. Hahn, W. Hong, T. Kirchhausen, P. Luzio, D. Root and R. Watanabe for providing DNA plasmids. This work was supported by the following grants to L.L.: NMRC/CBRG/007/2012, AcRF Tier1
RG132/15 and AcRF Tier1 RG48/13.

## Author contributions

L.L. and V.M. designed the study. V.M. performed experiments. V.M. and L.L. analyzed data. L.L. wrote the manuscript.

## Competing interests

The authors declare that they have no competing financial interests.

## Supplementary Information

### Supplemental Materials and Methods

#### DNA plasmids

The molecular cloning was conducted according to standard protocols. Constructs that were generated by PCR or oligonucleotide annealing have been confirmed by sequencing.

##### Vectors

To construct <u>pMyc-N1</u> vector, the oligonucleotides 5’-GAT CCA GTA CTC GAA CAA AAA CTC ATC TCA GAA GAG GAT CTG TAA AGC-3’ and 5’-GGC CGC TTT ACA GAT CCT CTT CTG AGA TGA GTT TTT GTT CGA GTA CTG-3’ were annealed and ligated into BamHI/NotI digested pEGFP-N1 vector (Clontech). To construct <u>pET30ax</u> vector, the oligonucleotides 5’-AGAT CCG AGA TCT GCT AGC GAA TTC CCG CGG GGA TCC C-3’ and 5’-GC TCT AGA CGA TCG CTT AAG GGC GCC CCT AGG GTT AA-3’ were annealed and ligated into BglII/EcoRI digested pET30a vector (Novagen). To make <u>pEGFP×2-N1</u> vector, the coding sequence (CDS) of GFP was amplified by PCR using pEGFP-N1 vector as the template and oligonucleotides 5’-GCA GTC GTC GAC GAT GGT GAG CAA GGG CGA G-3’ and 5’-GAC TGC GGA TCC TGC TTG TAC AGC TCG TCC AT-3’. The resulting PCR product was ligated into pEGFP-N1 vector using SalI/BamHI sites. To make <u>pEGFP×3-N1</u> vector, the CDS of GFP was amplified by PCR using oligonucleotides 5’-GCA GTC GAA TTC AGA TGG TGA GCA AGG GCG AG-3’ and 5’-GAC TGC GTC GAC TGC TTG TAC AGC TCG TCC AT-3’ and ligated into pEGFP×2-N1 vector using EcoRI/SalI sites. To clone construct <u>pHA-N1</u> vector, the oligonucleotides 5’-GaT CCA TAC CCA TAC GAC GTG CCA GAC TAC GCA TAA GC-3’ and 5’-GGC CGC TTA TGC GTA GTC TGG CAC GTC GTA TGG GTA TG-3’ for HA tag were annealed and ligated into BamHI/NotI digested pEGFP-N1 vector.

##### CD8a chimeras

To clone <u>CD8a</u> in pCI-neo vector (Promega), the CDS of CD8a (Mahajan et al., 2013) was PCR amplified using oligonucleotides 5’-GGA GCT GAA TTC GCC ACC ATG GCC TTA CCA GTG ACC GCC-3’ and 5’-GCT TCG TGG ATC CGT GAC GTA TCT CGC CGA AAG GCT G-3’ as primers and ligated into EcoRI/XbaI digested pCI-neo vector using the same sites. To construct <u>CD8a-GFP</u>, the CDS of CD8a was PCR amplified using oligonucleotides 5’-GCA GTC CTC GAG ATG GCC TTA CCA GTG ACC-3’ and 5’-CAC GTC GAA TTC GGA CGT ATC TCG CCG AAA GGC-3’ as primers and ligated into pEGFP-N1 vector using XhoI/EcoRI sites. <u>CD8a-sortilin</u> was previously described (Mahajan et al., 2013). To clone construct <u>CD8a-GFP×2 and -GFP×3</u>, the CDS of CD8a were released by digesting CD8a-GFP construct using XhoI/EcoRI and ligated into pEGFP×2-N1 and pEGFP×3-N1 vector respectively using the same sites. To clone CD8aAcyto-GFP (an intermediate clone), the extracellular and transmembrane domain of CD8a was PCR amplified using oligonucleotides 5’-GCA GTC CTC GAG ATG GCC TTA CCA GTG ACC-3’ and 5’-CCA TCT GAA TTC G GGT GAT AAC CAG TGA CAG-3’ as primers and ligated into pEGFP-N1 vector using XhoI/EcoRI sites. To clone construct <u>CD8aΔcyto-HA</u>, the CDS of CD8a was released by digesting CD8aAcyto-GFP with XhoI/EcoRI and ligated into pHA-N1 vector using the same sites. To construct <u>CD8a-cNLS-GFP</u>, the CDS of CD8a was PCR amplified using CD8a-GFP as the template and oligonucleotides 5’-GCA GTC CTC GAG ATG GCC TTA CCA GTG ACC-3’ and 5’-GAA TTC CTA CCT TTC TCT TCT TTT TTG GAT CTA CCT TTC TCT TCT TTT TTG GAT CGA CGT ATC TCG CCG AAA GG-3’ as primers. The resulting fragment was digested by XhoI/EcoRI and ligated into pEGFP-N1 vector using the same sites. To construct <u>CD8a-cNLS-GFP×2 and -GFP×3</u>, CD8a-cNLS was released by digesting CD8a-cNLS-GFP with XhoI/EcoRI and ligated into pEGFP×2-N1 and pEGFP×3-N1 vector respectively using the same sites. To construct <u>IBB-GFP</u>, the CDS of IBB was PCR amplified using IBB-tdTomato (Lu et al., 2011) as the template and 5’-GAT GCA GAA TTC GCC ACC ATG GCC GAG AAC CCC AGC TTG G-3’ and 5’-CAT GTT GGA TCC CGT GAA TCT TCT AGA CTT TCT TCT TGG-3’ as primers. The resulting product was digested by EcoRI/BamHI and ligated into pEGFP-N1 vector using the same sites. To construct <u>CD8a-IBB</u>, two PCR amplifications were performed by using CD8a and IBB-GFP as templates and primer pairs (5’-GcA GTC CTC GAG ATG gCc TTA CCA GTG ACC-3’ and 5’-CAA GCT GGG GTT CTC GGC GAC GTA TCT CGC CGA AAG-3’) and (5’-CTT TCG GCG AGA TAC GTC GCC GAG AAC CCC AGC TTG-3’ and 5’-CAC GTC GAA TTC CTA TTC TAG ACT TTC TTC TTG GG-3’) respectively. The two PCR fragments were mixed and subjected to the second round of PCR amplification using primers 5’-GCA GTC CTC GAG ATG GCC TTA CCA GTG ACC-3’ and 5’-CAC GTC GAA TTC CTA TTC TAG ACT TTC TTC TTG GG-3’. The resulting PCR product was digested by XhoI/EcoRI and ligated into pCI-neo vector using the same sites. To construct <u>CD8a-IBB-GFP</u>, the CDS of CD8a-IBB was PCR amplified using CD8a-IBB as the template and oligonucleotides 5’-GCA GTC CTC GAG ATG GCC TTA CCA GTG ACC-3’ and 5’-CAC GTC GGA TCC GC TTC TAG ACT TTC TTC TTG GG-3’ as primers. The resulting fragment was digested by XhoI/EcoRI and ligated into pEGFP-N1 vector using the same sites. To construct <u>CD8a-IBB-GFP×2 and -GFP×3</u>, the CDS of CD8a-IBB was released by digesting CD8a-IBB-GFP with XhoI/EcoRI and ligated into pEGFP×2-N1 and pEGFP×3-N1 vector respectively using the same sites. To construct <u>CD8a-hnRNP-M-GFP</u>, the CDS of CD8a was successively PCR amplified three times using CD8a-GFP as the template and the forward primer 5’-GCA GTC CTC Gag ATG GCC TTA CCA GTG ACC-3’ sequentially in combination with reverse primers 5’-CTC CTT CCT CTT CTC GTT CTG GGC GGG CCT CTC CAT GAC GTA TCT CGC CGA AAG-3’, 5’-GGG CTC GAA CCT GTT GCC GCC CCT CTT GAT GTT CTT CTC CTT CCT CTT CTC GTT-3’ and 5’-ACT GCA GAA TTC CCC TCT TGG TGG GGT TGG CGT AGG GCT CGA ACC TGT TGC C-3’. The resulting PCR product was digested by XhoI/EcoRI and ligated into pEGFP-N1 vector using the same sites. To construct <u>CD8a-hnRNP-M-GFP×3</u>, the CDS of CD8a-hnRNP-M was released by digesting CD8a-hnRNP-M-GFP with XhoI/EcoRI and ligated into pEGFP×3-N1 vector using the same sites. To construct <u>CD8a-CD4-GFP</u>, two PCR amplifications were performed by using CD8a and CD4 (Genbank: BC025782) as templates and primers pairs (5’-GCA GTC CTC GAG ATG GCC TTA CCA GTG ACC and CAG CAC AAT CAG GGC CAT ATC ACA GGC GAA GTC CAG-3’) and (5’-CTG GAC TTC GCC TGT GAT ATG GCC CTG ATT GTG CTG and CGA TCG GAA TTC TAA TGG GGC TAC ATG TCT TCT G-3’) respectively. The two PCR fragments were mixed and subjected to the second round of PCR amplification using primers (5’-GCA GTC CTC GAG ATG GCC TTA CCA GTG ACC and CGA TCG GAA TTC TAA TGG GGC TAC ATG TCT TCT G-3’). The resulting PCR product was digested by XhoI/EcoRI and ligated into pEGFP-N1 vector using the same sites.

##### Constructs of PM proteins

To construct <u>IL2Rα-GFP</u>, the CDS of IL2Rα was amplified by PCR using IL2Rα/hE-cadherin-cytotail (Addgene #45773; a gift from B. Gumbiner) as the template and primers 5’-GCA GTC CTC GAG ATG GAT TCA TAC CTG CTG ATG TGG-3’ and 5’-CAC TAC GAA TTC GGA TTG TTC TTC TAC TCT TCC TCT GTC TCC GCT GCC AGG TGA GCC CAC TCA GGA G-3’. The resulting PCR product was digested by NheI/EcoRI and ligated into pEGFP-N1 vector using the same sites. To construct <u>GFP-CAAX</u>, oligonucleotides 5’-AAT TCA GGC TGC ATG AGC TGC AAG TGT GTG CTC TCC TGA G-3’ and 5’-GAT CCT CAG GAG AGC ACA CAC TTG CAG CTC ATG CAG CCT G-3’ were annealed and ligated into EcoRI/BamHI digested pEGFP-C1 vector. To construct <u>Vamp2-mut-GFP</u> which has V43A and M46A mutations in Vamp2, two PCR amplifications were performed by using Vamp2-GFP (a gift from T. Galli) as the template and primer pairs (5’-TCA GAT CTC GAG ATG TCG GCT ACC GCT GCC and CCT GGC GAT GTC CGC CAC CTC ATC CAC CTG GGC CTG-3’) and (5’-GTG GCG GAC ATC GCC AGG GTG AAT GTG GAC AAG and ACT GCA GAA TTC GAG AGC TGA AGT AAA CTA TG-3’). The two PCR fragments were mixed and subjected to the second round of PCR amplification using primers 5’-TCA GAT CTC GAG ATG TCG GCT ACC GCT GCC-3’ and 5’-ACT GCA GAA TTC GAG AGC TGA AGT AAA CTA TG-3’. The resulting PCR product was digested by XhoI/EcoRI and ligated into pEGFP-N1 vector using the same sites. To construct <u>Vamp8-mut-GFP</u> which has V23P, K24A and M27A mutations in Vamp8, two PCR amplifications were performed by using Vamp8-GFP (a gift from P. Luzio) as the template and primer pairs (5’-TCA GAT CTC GAG ATG GAG GAG GCC AGT GGG and GGT GGC AAT ATT GGC GGG TCC CTC CAC CTC ACT CTG-3’) and (5’-CCC GCC AAT ATT GCC ACC CAG AAT GTG GAG CGG-3’ and 5’-ACT GCA GAA TTC GAG TGG GGA TGG TAC CAG TG-3’). The two PCR fragments were mixed and subjected to the second round of PCR amplification using primers 5’-TCA GAT CTC GAG ATG GAG GAG GCC AGT GGG-3’ and 5’-ACT GCA GAA TTC GAG TGG GGA TGG TAC CAG TG-3’. The resulting PCR product was digested by XhoI/EcoRI and ligated into pEGFP-N1 vector using the same sites.

##### Fibrocystin constructs

To construct <u>CFFAC-GFP</u>, the extracellular domain of CD8a was successively PCR amplified three times using CD8a-GFP as the template and the forward primer 5’-GCA GTC CTC GAG ATG GCC TTA CCA GTG ACC-3’ sequentially in combination with reverse primers 5’-CCA CGC TGC ACA GGC TCA GGG CCA GGA TGA TGG TGC TTC TCT CAT CAC AGG CGA AGT CC-3’, 5’-CTT AAA CCA GCA GCA AAC GAG GCA GCT CAG GGC CAC CCA GCT GGC CAC GCT GCA CAG GC-3’ and 5’-GCT CAT GAA TTC GTT CTG GCT TTA TCT TTC TGG TCT TGC TCT TCT TAA ACC AGC AGC A-3’. The resulting PCR product was digested by XhoI/EcoRI and ligated into pEGFP-N1 vector using the same sites. To construct <u>CFFΔC-Myc</u>, the CDS of CFFΔC was released by digesting CFFAC-GFP with XhoI/EcoRI and ligated into pMyc-N1 vector using the same sites. To clone <u>CFF-GFP</u>, the extracellular domain of CD8a and transmembrane domain of fibrocystin was PCR amplified using CFFAC-GFP as the template and the following oligonucleotides as primers 5’-GCA GTC CTC GAG ATG GCC TTA CCA GTG ACC-3’ (primer F) and 5’-CCA AGA TGA TAG TTG ATC TCT CAT CAC AGG CGA AGT CCA G-3’. The cytosolic domain of fibrocystin was PCR amplified using IMAGE clone (GenBank Accession No.:4238864) as the template and the following oligonucleotides as primers: 5’-CTG GAC TTC GCC TGT GAT GAG AGA TCA ACT ATC ATC TTG G-3’ and 5’-ACT GCA GAA TTC GCT GGA TGG TTT CTG GTG G-3’ (primer R). The mixture of two PCR fragments were subjected to PCR amplification by primer F and R and the resulting fragment was digested by XhoI/EcoRI and ligated into pEGFP-N1 vector using the same sites. To construct <u>CFF-Myc and -HA</u>, the CDS of CFF was released by digesting CFF-GFP with XhoI/EcoRI and ligated into pMyc-N1 and pHA-N1 vector using the same sites respectively. To construct <u>CD8a-f-CTS-</u> <u>GFP</u> the CDS of CD8a-f-CTS was PCR amplified using primer pair (5’-GTC TAG AAT TCA GCC ACC ATG GCC TTA CCA GTG ACC GCC TTG C-3’ and 5’-AAT TCG TTC TGG CTT TAT CTT TCT GGT CTT GCT CTT CTT AAA CCA GCA GCA AAC GAG GCA CAT C-3’). The resulting PCR product was digested by EcoRI/BamHI and ligated into pEGFP-N3 vector (Clontech) using the same sites. To construct <u>f-CTS-</u> <u>GFP</u>, two oligonucleotides 5’-AAT TCA TGT GCC TCG TTT GCT GCT GGT TTA AGA AGA GCA AGA CCA GAA AGA TAA AGC CAG AAC GG-3’ and 5’-GAT CCC GTT CTG GCT TTA TCT TTC TGG TCT TGC TCT TCT TAA ACC AGC AGC AAA CGA GGC ACA TG-3’ were annealed and ligated into EcoRI/BamHI digested pEGFP-N1 vector. To construct GFP-f-CTS (an intermediate clone), the CDS of f-CTS and part of the 3’ non-coding region was PCR amplified using CD8a-f-CTS as the template and 5’-GCA GTC GAA TTC TGC CTC GTT TGC TGC TGG-3’ and 5’-CAC GTC GGA TCC CTG TTG GGA AGG GCG ATC GG-3’ as primers. The resulting PCR product was digested by EcoRI/BamHI and ligated into pEGFP-C2 vector (Clontech) using the same sites. To construct <u>GST-f-CTS</u>, the fragment containing f-CTS was released from GFP-f-CTS with EcoRI/BamHI and ligated into pGEB vector, a modified pGEX-KG vector (GE Healthcare Life Sciences), using the same sites.

##### Constructs of ciliary residents

To construct <u>GST fused CTS of cystin, polycystin-1, polycystin-2, prRDH and pheripherin</u>, oligonucleotide pairs (5’-AA TTC ACC GCC AGC GAG GGC GGC ACC GCC CGG-3’ and 5’-GA TCC CGG GCG GTG CCG CCC TCG CTG GCG GTG-3’), (5’-AA TTC AAG GTG CAC CCC AGC AGC ACC CGG-3’ and 5’-GA TCC CGG GTG CTG CTG GGG TGC ACC TTG-3’), (5’-AA TTC ATG GTG AAC AGC AGC AGG GTG CAG CCC CAG CAG CCC GGC GAC GCC CGG-3’ and 5’-GA TCC CGG GCG TCG CCG GGC TGC TGG GGC TGC ACC CTG CTG CTG TTC ACC ATG-3’), (5’-AA TTC CTG AGG TGC CTG GCC TGC AGC TGC TTC AGG ACC CCC GTG TGG CCC AGG CGG-3’ and 5’-GA TCC CGC CTG GGC CAC ACG GGG GTC CTG AAG CAG CTG CAG GCC AGG CAC CTC AGG-3’) and (5’-AATTC AAG AGC AGC TGG GAG CTG GTG AAG AGC ATG GGC AAG CTG AAC AAG GTG GAG ACC GCC GGC CGG-3’ and 5’-GAT CCC GGC CGG CGG TCT CCA CCT TGT TCA GCT TGC CCA TGC TCT TCA CCA GCT CCC AGC TGC TCT TG-3’) were annealed and ligated into EcoRI/BamHI digested pGEB vector respectively. To construct <u>GST-rhodopsin-CTS</u>, the c-terminal tail of rhodopsin was PCR amplified using rhodopsin-GFP (Addgene #45399; a gift from D. Williams) as the template and oligonucleotides 5’-CTC GAG AAT TCA TGG GCT GCT TCT TCT CC-3’ and 5’-CAG TAA GGA TCC TTA GGC AGG CGC CAC TTG-3’ as primers. The resulting PCR product was digested by EcoRI/BamHI and ligated into pGEB vector using the same sites. To construct <u>GST-RP2</u>, the CDS of RP2 was PCR amplified using Addgene plasmid #23636 (a gift from W. Hahn and D. Root) as the template and oligonucleotides 5’-GCA TCA GAA TTC ATG AAC AAG CAG TTC CGG-3’ and 5’-CCC GGG GTC GAC TAT TCC CAT CTG TAT ATC-3’ as primers. The resulting PCR product was digested by EcoRI/SalI and ligated into pGEX-4T1 vector (GE healthcare) using the same sites. To construct <u>RP2-GFP</u>, the CDS of RP2 was released by digesting GST-RP2 with EcoRI/SalI and ligated into pEGFP-N3 vector (Clontech) using the same sites. To construct <u>GST-RP2-C86Y,P95L</u>, two PCR amplifications were performed by using GST-RP2 as the template and primer pairs (5’-CTC GAG AAT TCA TGG GCT GCT TCT TCT CC-3’ and 5’-GCC TTT CAC CAG TCC CAG AAA AAT TAT GCA GTT AGT GTA GTC ATC-3’) and (5’-GAT GAC TAC ACT AAC TGC ATA ATT TTT CTG GGA CTG GTG AAA GGC-3’ and 5’-CCC GGG GTC GAC ATT CCC ATC TGT ATA TC-3’). The two PCR fragments were mixed and subjected to the second round of PCR amplification using primers 5’-CTC GAG AAT TCA TGG GCT GCT TCT TCT CC-3’ and 5’-CCC GGG GTC GAC ATT CCC ATC TGT ATA TC-3’. The resulting PCR product was digested by EcoRI/SalI and ligated into pGEX-4T1 vector using the same sites. To construct <u>SSTR3-GFP</u>, the CDS of SSTR3 was PCR amplified from IMAGE clone 100000442 (GenBank Accession No.:NM_001051.2) using primers 5’-TAC CtC GAG ATG GAC ATG CTT CAT CC-3’ and 5’-CGT AAG CTT CAG GTA GCT GAT GCG CAT-3’. The resulting PCR product was digested by XhoI/HindIII and ligated into pEGFP-N1 vector using the same sites. To construct <u>GST-SSTR3</u>, the CDS of SSTR3i3 (Jin et al., 2010) was PCR amplified from SSTR3-GFP using primers 5’-GCA GTC GAA TTC ATG GTG GTG AAG GTG CGC TCA G-3’ and 5’-ACG ACT GGA TCC TTA GCG CGT GAC CCT GCG TTC-3’. The resulting PCR product was digested by EcoRI/BamHI and ligated into pGEB vector using the same sites. To construct <u>CD8a-prRDH-CTS-GFP</u>, oligonucleotides 5’-AAT TCC TGA GGT GCC TGG CCT GCA GCT GCT TCA GGA CCC CCG TGT GGC CCA GGC GG-3’ and 5’-GAT CCC GCC TGG GCC ACA CGG GGG TCC TGA AGC AGC TGC AGG CCA GGC ACC TCA GG-3’ were annealed and ligated into CD8a-GFP using EcoRI/BamHI sites.

##### Constructs of Rab8

To construct <u>GFP-Rab8-wt</u>, the CDS of Rab8a was PCR amplified using IMAGE clone 3547214 (GenBank Accession No.: BC002977.1) as the template and oligonucleotides 5’-GTC TAG GAA TTC GCG AAG ACC TAC GAT TAC CTG TTC AAG-3’ and 5’-CAC TAC GGA TCC CTA CAG AAG AAC ACA TCG GAA AAA GCT GCT-3’. The resulting PCR product was digested by EcoRI/BamHI and ligated into pEGFP-C2 vector using the same sites. To construct <u>GFP-Rab8-T22N</u>, two PCR amplifications were performed by using GFP-Rab8-wt as the template and primer pairs (5’-GTC TAG GAA TTC GCG AAG ACC TAC GAT TAC CTG TTC AAG-3’ and 5’-GGA ACA GGA CAC AGT TCT TCC CCA CCC C-3’) and (5’-GGG GTG GGG AAG AAC TGT GTC CTG TTC C-3’ and 5’-CAC TAC GGA TCC CTA CAG AAG AAC ACA TCG GAA AAA GCT GCT-3’). The two PCR fragments were mixed and subjected to the second round of PCR amplification using primers 5’-GTC TAG GAA TTC GCG AAG ACC TAC GAT TAC CTG TTC AAG-3’ and 5’-CAC TAC GGA TCC CTA CAG AAG AAC ACA TCG GAA AAA GCT GCT-3’. The resulting PCR product was digested by EcoRI/BamHI and ligated into pEGFP-C2 vector using the same sites. <u>GFP-Rab8-Q67L</u> was similarly constructed in pEGFP-C2 vector using primer pairs (5’-GTC TAG GAA TTC GCG AAG ACC TAC GAT TAC CTG TTC AAG-3’ and 5’-CCG AAA CCG TTC CAG ACC GGC TGT GTC-3’) and (5’-GAC ACA GCC GGT CTG GAA CGG TTT CGG-3’ and 5’-CAC TAC GGA TCC CTA CAG AAG AAC ACA TCG GAA AAA GCT GCT-3’). To construct <u>GST-or His-Rab8-wt, -T22N</u> <u>and -Q71L</u>, CDSs of Rab8 were released by digesting GFP-Rab8-wt, -T22N and -Q67L with EcoRI/BamHI respectively and ligated into pGEB or pET30ax vector using the same sites. To construct <u>Myc-Rab8-wt, -T22N and -Q67L</u>, CDSs of Rab8 were released by digesting GFP-Rab8-wt, -T22N and -Q67L with EcoRI/BamHI respectively and ligated into a pCI-neo vector containing a double-Myc-tag using the same sites.

##### Constructs of Arl13b

To construct <u>Arl13b-GFP</u>, the CDS of Arl13b was amplified by PCR from IMAGE clone 40026399 (GenBank Accession No.: BC104036) using primers 5’-GTA ATG CTC GAG GCC ACC ATG TTC AGT CTG ATG GCC AGT T-3’ and 5’-CTA CGG ATC CAA TGA GAT CAC ATC ATG AGC ATC AC-3’. The resulting PCR product was digested by XhoI/BamHI and ligated into pEGFP-N1 vector using the same sites. To construct <u>Arl13b-mCherry</u>, the CDS of Arl13b was released by digesting Arl13b-GFP with XhoI/BamHI and ligated into pmCherry-N1 (Clontech) using the same sites. To construct <u>His-Arl13b-C-ter</u>, the CDS of Arl13b c-terminal fragment was PCR amplified using Arl13b-GFP as the template and primers 5’-GCAGTC AGATCT C GAATTC GAG TTG GAC CCA GAA CCA ACG-3’ and 5’-CACGTC CTCGAG GGATCC TTA TGA GAT CAC ATC ATG AGC ATC-3’. The resulting PCR product was digested by BglII/XhoI and ligated into pET30a vector using the same sites. To construct <u>GST-Arl13b-C-ter</u>, the CDS of Arl13b-C-ter was released by digesting His-Arl13b-C-ter with EcoRI/BamHI and ligated into pGEB vector using the same sites. To construct dyn-1(K44A)-Myc, the CDS of dyn-1(K44A) was released by digesting dyn-1(K44A)-GFP (Addgene #22197; a gift from Pietro De Camilli) with XhoI/BamHI and ligated into pMyc-N1 vector using the same sites.

##### Constructs of TNPO1

To construct <u>Myc-TNPO1</u>, the CDS of TNPO1 was PCR amplified from the cDNA clone SC109373 (Origene; GenBank Accession No.: NM_153188) using primers 5’-TCA GAT CTC GAG ATG GAG TAT GAG TGG AAA CC-3’ and 5’-TCC GGT GGA TCC TTA AAC ACC ATA AAA AGC TG-3’. The resulting PCR product was digested by XhoI/BamHI and ligated into pDMyc-neo vector using the same sites. To construct TNPO1 <u>Myc-Δ315, -Δ539 and -Δ699</u>, PCR amplifications were conducted using Myc-TNPO1 as the template and forward primer 5’-CGT TGA CTC GAG ATG AAG TAC TCA GAC ATA G-3’, 5’-CGT TGA CTC GAG ATG TAC CAG CAT AAG AAC CTG-3’ and 5’-CGT TGA CTC GAG ATG CCA ATA TTG GGA ACC-3’, respectively, in combination with reverse primer 5’-TCC GGT GGA TCC TTA AAC ACC ATA AAA AGC TG-3’. pCr products were digested by XhoI/BamHI and ligated into pDMyc-neo vector using the same sites.

##### Constructs of Ran GTPase

<u>mCherry-Ran-Q69L</u> was from Addgene (# 30309; a gift from J. Brenman). To construct <u>GST-Ran-wt</u>, two PCR amplifications were performed by using mCherry-Ran-Q69L as the template and primer pairs (5’-TCT AGC GAA TTC ATG GCT GCG CAG GGA GAG-3’ and 5’-CCA CCG AAT TTC TCC TGG CCG GCT GTG TCC C-3’) and (5’-GGG ACA CAG CCG GCC AGG AGA AAT TCG GTG G-3’ and 5’-TAG CAG GGA TCC TCA CAG GtC ATC ATC CTC ATC-3’). The two fragments were mixed and subjected to the second round of PCR amplification using primers 5’-TCT AGC GAA TTC ATG GCT GCG CAG GGA GAG-3’ and 5’-TAG CAG GGA TCC TCA CAG GTC ATC ATC CTC ATC-3’. The resulting PCR product was digested by EcoRI/BamHI and ligated into pGEB vector using the same sites. To construct <u>GST-Ran-</u> <u>Q69L</u>, the CDS of Ran-Q69L was PCR amplified using mCherry-Ran-Q69L as the template and 5’-TCT AGC GAA TTC ATG GCT GCG CAG GGA GAG-3’ and 5’-TAG CAG GGA TCC TCA CAG GTC ATC ATC CTC ATC-3’ as primers. The resulting PCR product was digested by EcoRI/BamHI and ligated into pGEB vector using the same sites. To construct <u>GST-Ran-T24N</u>, two PCR amplifications were performed by using GST-Ran-wt as the template and primer pairs (5’-TCT AGC GAA TTC ATG GCT GcG CAG GGA GAG-3’ and 5’-CGT TTC ACG AAG GTG TTT TTT CCA GTA CCA CC-3’) and (5’-GGT GGT ACT GGA AAA AAC ACC TTC GTG AAA CG-3’ and 5’-TAG CAG GGA TCC TCA CAG GTC ATC ATC CTC ATC-3’). The two PCR fragments were mixed and subjected to the second round of PCR amplification using primers 5’-TCT AGC GAA TTC ATG GCT GCG CAG GGA GAG-3’ and 5’-TAG CAG GGA TCC TCA CaG GTC ATC ATC CTC ATC-3’. The resulting PCR product was digested by EcoRI/BamHI and ligated into pGEB vector using the same sites.

##### shRNAs

To construct lentiviral shRNA targeting TNPO1, oligonucleotides 5’-CCGG CAA TGC TCA ACC AGA TCA ATA CTC GAG TAT TGA TCT GGT TGA GCA TTG TTT TTG-3’ and 5’-AATTC AAAAA CAA TGC TCA ACC AGA TCA ATA CTC GAG TAT TGA TCT GGT TGA GCA TTG-3’ were annealed and ligated into AgeI/EcoRI digested pLKO.1-TRC vector (Addgene # 10878; a gift from D. Root). Lentiviral shRNA targeting GL2 (negative control) was similarly constructed using oligonucleotides 5’-CCGG AAC GTA CGC GGA ATA CTT CGA CTC GAG TCG AAG TAT TCC GCG TAC GTT TTT TTG-3’ and 5’-AATTC AAAAA AAC GTA CGC GGA ATA CTT CGA CTC GAG TCG AAG TAT TCC GCG TAC GTT-3’.

The following plasmids were generous gifts: Vamp5-GFP from W. Hong, mCherry-clathrin light chain from T. Kirchhausen and mCherry-CD59 from R. Watanabe (Addgene # 50378).

---

**Figure S1.**
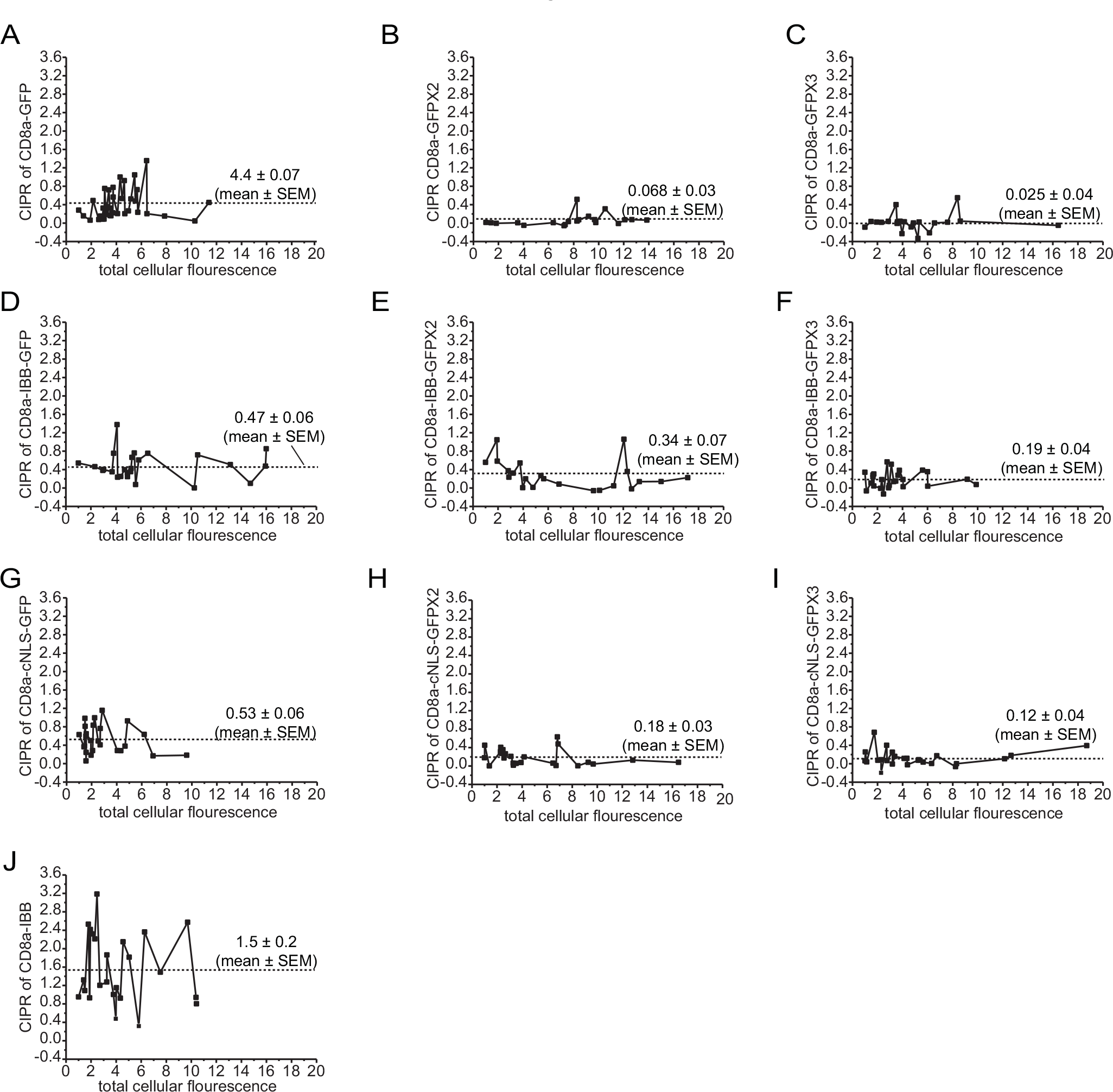
The CIPR of a membrane protein is independent of its expression level. (A-J) Ciliated RPE1 cells expressing different CD8a chimeras were analyzed for CPIRs and the total cellular intensity of CD8a chimeras. CPIRs are plotted against the total cellular intensity of CD8a chimera (arbitrary unit). Plots are organized similarly to Figure 1C. Data were from n=25 cells.

**Figure S2.**
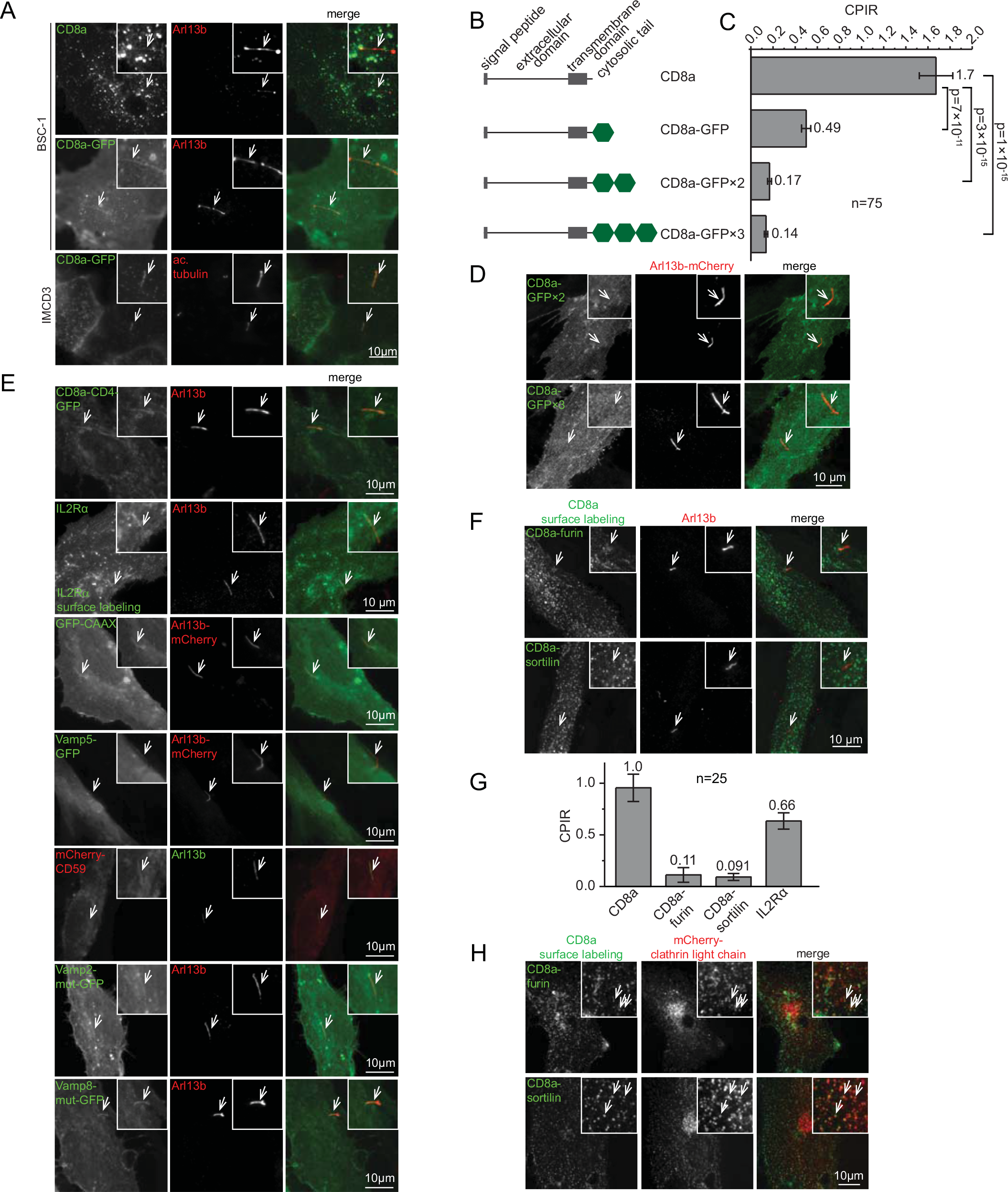
PM-localized membrane proteins can be found at cilia. Cells were transiently transfected by various DNA plasmids and subsequently subjected to induction for ciliogenesis. (A) The ciliary localization of CD8a or CD8a-GFP in BSC-1 or IMCD3 cells. (B) A schematic diagram showing CD8a chimeras tagged by 0-3 copies of GFP. (C) The ciliary localization of CD8a chimera is dependent on its cytosolic size. CPIRs of CD8a chimeras tagged by 0-3 copies of GFP. Error bar, SEM. n=75 cells. The mean value is indicated at the right of each column. p values (t-test) of selected pairs are denoted. (D) Images of CD8a-GFP×2 and CD8a-GFP×3 showing that they are virtually devoid of ciliary localization. (E) CD8a-CD4-GFP, IL2Ra, GFP-CAAX, Vamp5-GFP, mCherry-CD59, Vamp2-mut-GFP, Vamp8-mut-GFP localized to cilia in ciliated RPE1 cells. (F) CD8a-furin and -sortilin did not localize to cilia. Cells expressing CD8a fusion chimeras were surface-labeled by anti-CD8a antibody. (G) CPIRs of CD8a, CD8a-furin, CD8a-sortilin and IL2Ra in ciliated RPE1 cells. Error bar, SEM. n=25 cells. The mean is indicated at the top of each column. The experiment was performed together with Figure 1E. Therefore, both share the same CPIR of CD8a. (H) CD8a-furin and -sortilin localized to clathrin-coated pits at the PM. RPE1 cells co-expressing mCherry-clathrin light chain and CD8a-furin or-sortilin were subjected to surface labeling by anti-CD8a antibody. Three examples of colocalization spots are indicated by arrows in each panel. For all panels, cilia were identified by endogenous Arl13b, acetylated tubulin (ac. tubulin) or co-expressed Arl13b-mCherry. In each image, an area of interest was enlarged and boxed at the upper right corner to show the positive or negative ciliary colocalization. The cilium of interest is indicated by an arrow. Scale bar, 10 μM.

**Figure S3.**
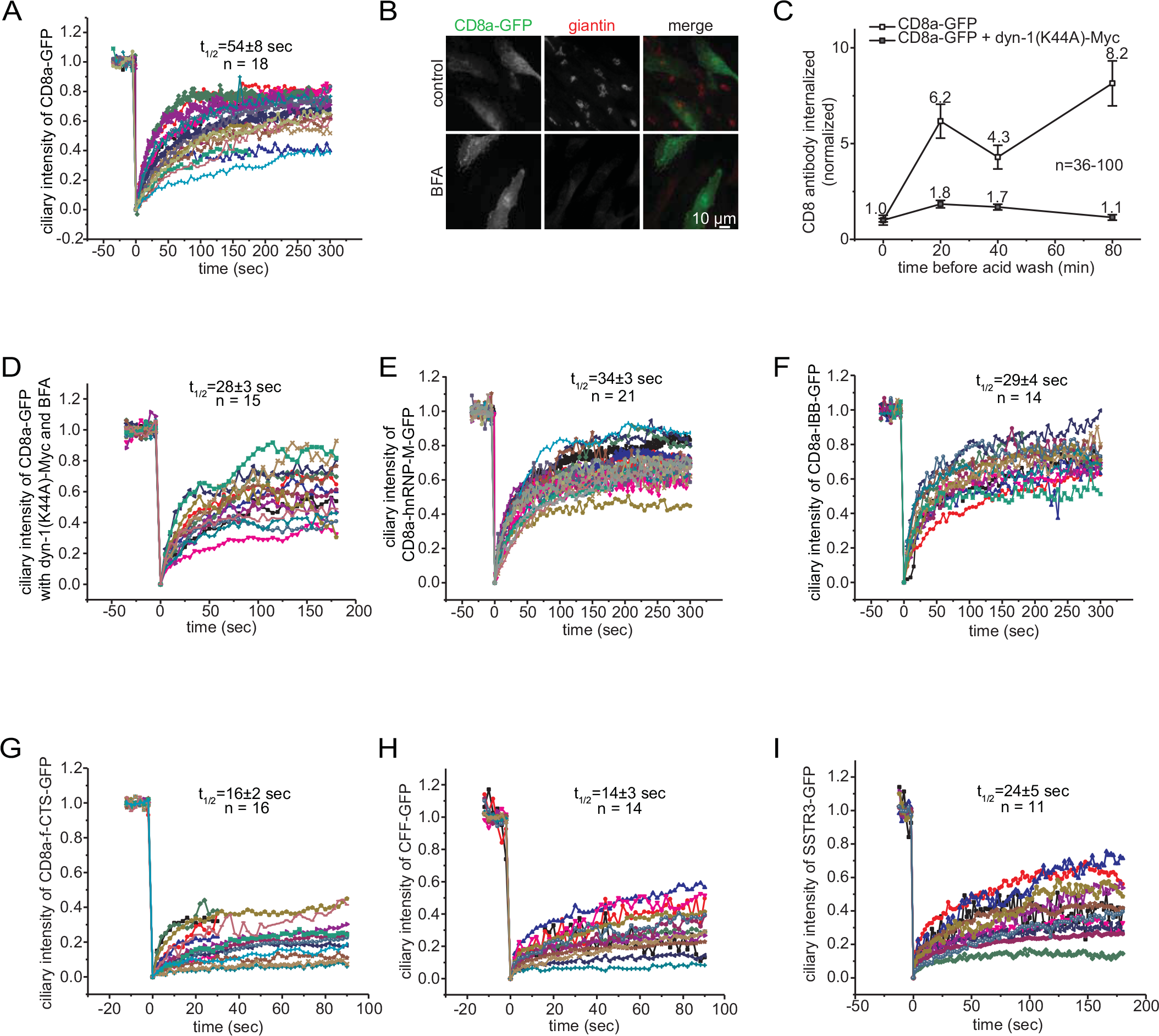
Whole cilium FRAP analysis of CD8a-GFP chimeras. (A) The intensity of CD8a-GFP at cilia is plotted against the time (sec) under control condition during whole cilium FRAP analysis. The experiment is described in Figure 1F. (B) RPE1 cells expressing CD8a-GFP were treated with 5 μM BFA or DMSO (control) for 30 min and processed for immunofluorescence labeling of endogenous giantin (a Golgi marker). Scale bar, 10 μm. (C) Dynamin-1(K44A) inhibited the endocytosis of CD8a-GFP. RPE1 cells expressing CD8a-GFP alone or together with dyn-1(K44A)-Myc were incubated with anti-CD8a antibody on ice for 60 min. After washing away unbound antibody, cells were warmed up to 37 °C for indicated time before treatment with ice cold acid to remove surface-bound antibody (acid wash). The internalized antibody was fluorescence stained. The total intensity of internalized antibody was divided by the total intensity of CD8a-GFP and plotted against the incubation time before acid wash. Error bar, SEM. Mean values are indicated. (D) The intensity of CD8a-GFP at cilia is plotted against time during whole cilium FRAP analysis in cells co-expressing dyn-1(K44A)-Myc and treated with BFA. The experiment corresponds to Figure 1F (E-I) Whole cilium FRAP traces. Experiments correspond to Figures 2E-G.

**Figure S4.**
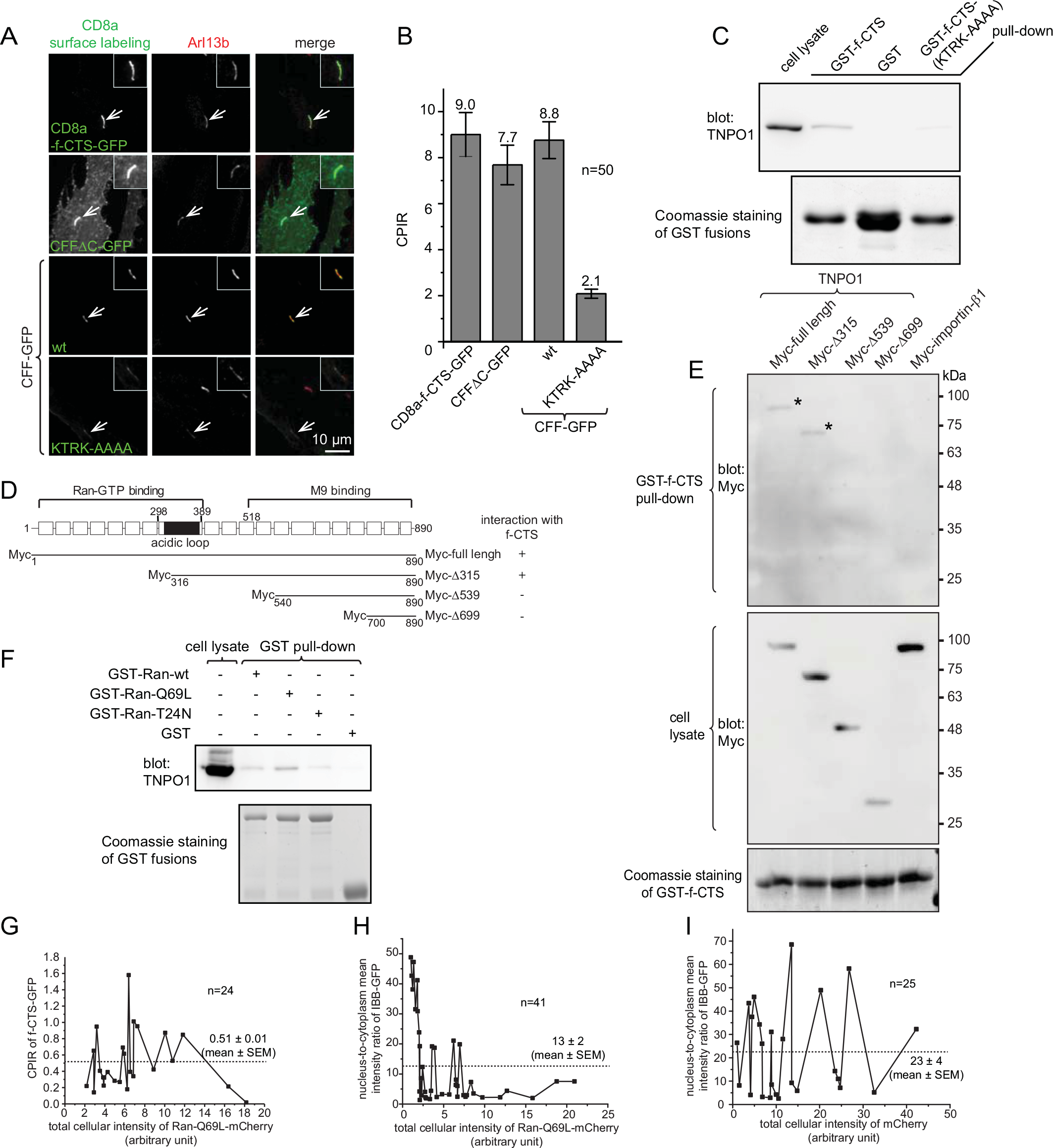
The interaction between f-CTS and TNPO1 is dependent on amino acids KTRK within f-CTS and the central region of TNPO1 but it is independent on Ran GTPase. (A and B) Images and CPIRs of CD8a-fused fibrocystin chimeras in ciliated RPE1 cells. The cilium of interest is indicated by an arrow, enlarged and boxed at the upper right corner of each image. To acquire the CPIR, CD8a chimeras were revealed by surface labeling. Scale bar, 10 μm. Error bar, SEM. n=50 cells. The mean value is indicated at the top of each column. (C) KTRK to AAAA mutation (KTRK-AAAA) greatly reduced the interaction between f-CTS and TNPO1. Bead-immobilized GST-f-CTS or GST-f-CTS-(KTRK-AAAA) were incubated with HEK293T cell lysate and the material pulled down was blotted for TNPO1. (D) A schematic diagram showing the domain organization and truncation clones of TNPO1 used in this study. (E) The interaction between f-CTS and TNPO1 requires the central region (amino acid 316-539) of TNPO1. Bead-immobilized GST-f-CTS was incubated with HEK293T cell lysate expressing Myc-tagged truncations of TNPO1 and the protein pulled down was blotted for Myc-tag. denotes the band for full-length or Myc-A315 protein. In selected gel blots, numbers at the right indicate the molecular weight markers in kDa. (F) Bead-immobilized GST-Ran-Q69L (GTP form mutant) pulled down significantly more endogenous TNPO1 from HEK293T cell lysate than corresponding T24N (GDP form mutant) and wild type GST-Ran. (G) The ciliary localization of f-CTS-GFP was not affected by the overexpression of Ran-Q69L-mCherry. Ciliated RPE1 cells co-expressing f-CTS-GFP and Ran-Q69L-mCherry were analyzed to plot the CPIR of f-CTS-GFP against the total cellular intensity of Ran-Q69L-mCherry. Data points are connected by lines from low to high total cellular intensity. n=24 cells. (H and I) The nucleus-to-cytoplasm mean intensity ratio of IBB-GFP decreased when the cellular amount of Ran-Q69L-mCherry increased, demonstrating the functionality of our Ran-Q69L-mCherry. RPE1 cells co-expressing IBB-GFP and Ran-Q69L-mCherry (H) or mCherry (I, as a negative control) were analyzed to plot the nucleus-to-cytoplasm mean intensity ratio of IBB-GFP against the total cellular intensity of Ran-Q69L-mCherry or mCherry. Data points are connected by lines from low to high total cellular intensity. n=41 (H) and 25 (I) cells.

**Figure S5.**
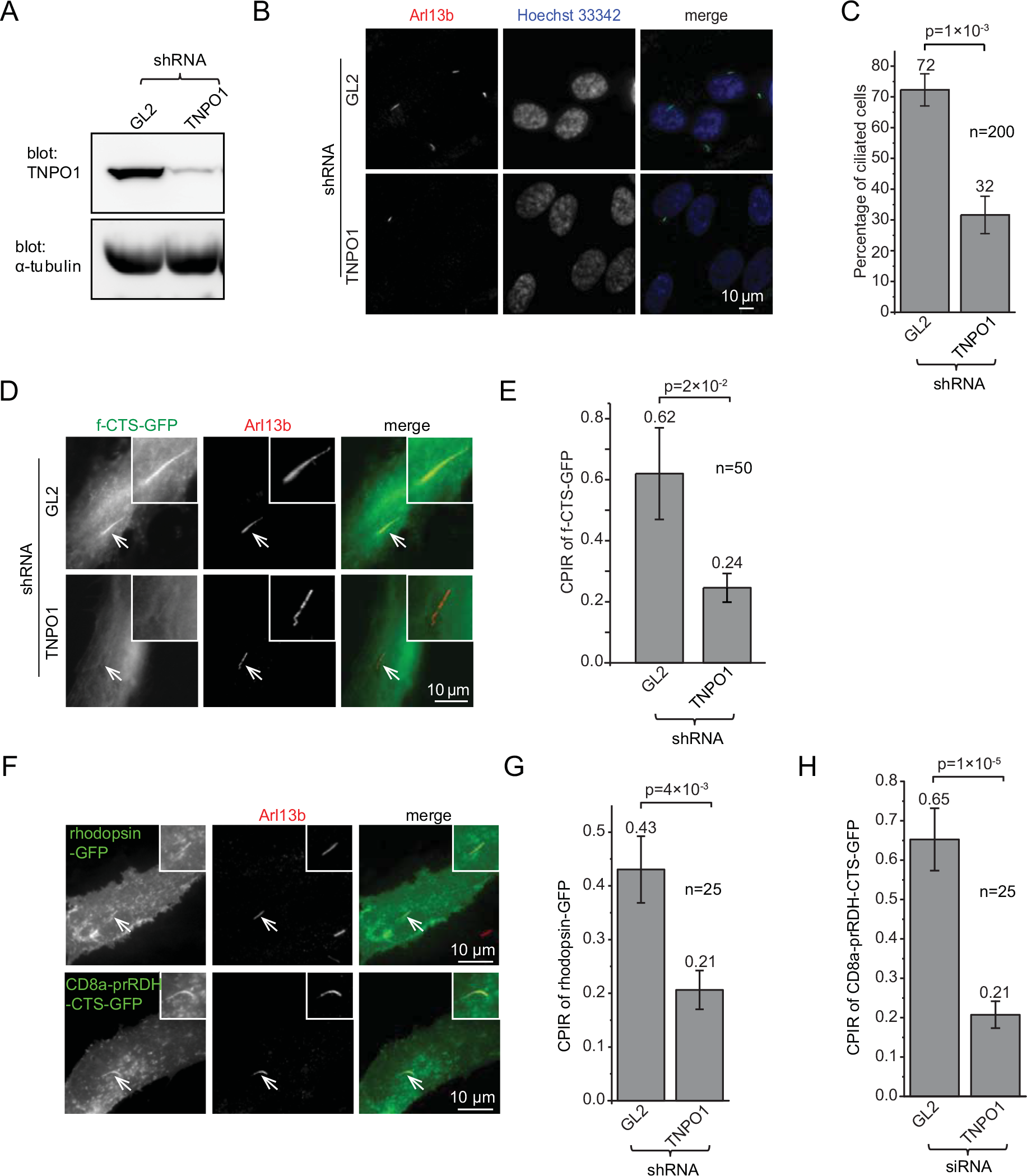
Depletion of TNPO1 reduces the ciliogenesis and ciliary localization of fibrocystin. (A) Endogenous TNPO1 was significantly depleted by shRNA-mediated knockdown of endogenous TNPO1. (B and C) The depletion of TNPO1 reduced the percentage of ciliated cells. TNPO1-depleted RPE1 cells were induced for ciliogenesis and ciliated cells were counted. Images and percentages are shown in (B) and (C) respectively. (D and E) TNPO1-depleted RPE1 cells were transfected to express f-CTS-GFP and induced for ciliogenesis. Images and CPIRs are shown in (D) and (E) respectively. (F) Full-length rhodopsin-GFP and CD8a-prRDH-CTS-GFP localized to cilia. (G and H) TNPO1-depleted RPE1 cells were transfected to express rhodopsin-GFP or CD8a-prRDH-CTS-GFP and induced for ciliogenesis. CPIRs are plotted. In (D and F), the cilium of interest is indicated by an arrow, enlarged and boxed at the upper right corner of each image. Scale bar, 10 μm. In (C, E, G and H), p values (t-test) between GL2 and TNPO1 and the number of cells, n, are indicated. The mean value is denoted at the top of each column. Error bar, SEM.

**Figure S6.**
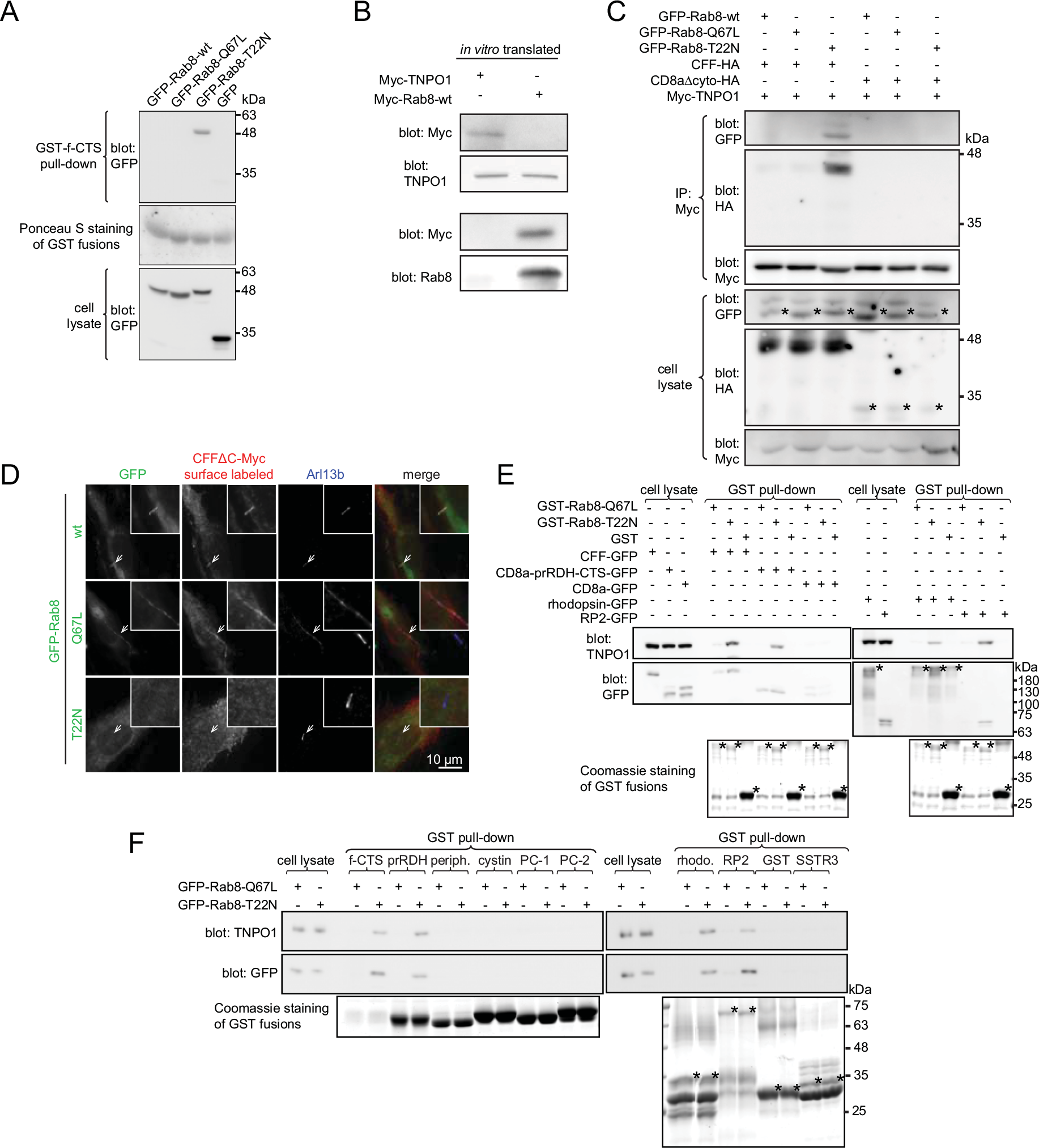
The interaction among CTSs, Rab8 and TNPO1. (A) Bead-immobilized GST-f-CTS specifically pulled down GFP-Rab8-T22N but not -Q67L and -wt from the cell lysate expressing respective GFP chimeras. (B) Rabbit reticulocyte lysate from in vitro transcription and translation system contained endogenous TNPO1 but not Rab8. Rabbit reticulocyte lysate was mixed with methionine and Myc-TNPO1 or Myc-Rab8 expression DNA plasmids that contain T7 promoters for 1 hour at 30 °C. The lysate was subsequently separated by electrophoresis and blotted for Myc-tag, TNPO1 and Rab8. The Rab8 antibody used here should cross-react with rabbit Rab8 as the identity of antigen regions between human and rabbit is as high as 97%. (C) TNPO1 indirectly interacted with Rab8-T22N via f-CTS. The experiment also provided further evidence showing that Rab8-T22N increases the interaction between TNPO1 and f-CTS. HEK293T cell lysate triply co-expressing Myc-TNPO1, one of the GFP-Rab8 mutants (wt, Q67L and T22N) and CFF-HA or CD8aAcyto-HA was subjected to IP using anti-Myc antibody and co-IPed proteins were blotted for HA-tag and GFP. HA was blotted by HRP conjugated anti-HA antibody while GFP was blotted by anti-GFP primary antibody followed by HRP conjugated protein A. (D) The effect of Rab8 guanine nucleotide binding mutants on the ciliary localization of fibrocystin. Ciliated RPE1 cells co-expressing CFFAC-Myc and GFP, GFP-Rab8-wt, -Q67L or -T22N were subjected to surface labeling by CD8a antibody to reveal the ciliary localization of CFFAC-Myc. Cilia were identified by endogenous Arl13b staining. The cilium of interest is indicated by an arrow, enlarged and boxed at the upper right corner of each image. Scale bar, 10 μm. (E) RP2, rhodopsin or the CTS of prRDH assembled a ternary complex with Rab8-T22N and TNPO1. Bead-immobilized GST or GST-Rab8 mutants were incubated with HEK293T cell lysate expressing CFF-GFP (positive control), CD8a-prRDH-CTS-GFP, CD8a-GFP (negative control), rhodopsin-GFP or RP2-GFP. The bound GFP fusion proteins and endogenous TNPO1 were subsequently blotted. Note the high apparent molecular weight of full-length rhodopsin-GFP in our experimental condition (marked by “*”). (F) The screen showing that RP2 and CTSs of fibrocystin, prRDH and rhodopsin, but not peripherin, cystin, polycystin-1, polycystin-2, and SSTR3, demonstrate enhanced interaction with endogenous TNPO1 in the present of Rab8-GDP mutant. GST-CTSs immobilized on beads were incubated with HEK293T cell lysate expressing GFP-Rab8-Q67L or -T22N. The bound GFP-Rab8 and endogenous TNPO1 were subsequently blotted. CTSs of fibrocystin and prRDH were positive controls. Periph., peripherin. PC-1, polycystin-1. PC-2, polycystin-2. rhodo., rhodopsin. “*” denotes the specific band. In selected gel blots, numbers at the right indicate the molecular weight markers in kDa.

